# Is there a latitudinal diversity gradient for symbiotic microbes? A case study with sensitive partridge peas

**DOI:** 10.1101/2023.05.03.539300

**Authors:** Tia L. Harrison, Zoe A. Parshuram, Megan E. Frederickson, John R. Stinchcombe

## Abstract

Mutualism is more prevalent in the tropics than temperate zones and is therefore expected to play an important role in generating and maintaining high species richness found at lower latitudes. However, results on the impact of mutualism on latitudinal diversity gradients are mixed, and few empirical studies sample both temperate and tropical regions. We investigated whether a latitudinal diversity gradient exists in the symbiotic microbial community associated with the legume *Chamaecrista nictitans*. We sampled bacteria DNA from nodules and the surrounding soil of plant roots across a latitudinal gradient (38.64 °N to 8.68 °N). Using 16S rRNA sequence data, we identified many non-rhizobial species within *C. nictitans* nodules that cannot form nodules or fix nitrogen. Species richness increased towards lower latitudes in the non-rhizobial portion of the nodule community but not in the rhizobial community. The microbe community in the soil did not predict the non-rhizobia community inside nodules, indicating that host selection is important for structuring non-rhizobia communities in nodules. We next factorially manipulated the presence of three non-rhizobia strains in greenhouse experiments and found that co-inoculations of non-rhizobia strains with rhizobia had a marginal effect on nodule number and no effect on plant growth. Our results suggest that these non-rhizobia bacteria are likely commensals – species that benefit from associating with a host but are neutral for host fitness. Overall, our study suggests that temperate *C. nictitans* plants are more selective in their associations with the non-rhizobia community, potentially due to differences in soil nitrogen across latitude.

## Introduction

How mutualisms influence latitudinal patterns of species richness (and vice versa) remains an open question in ecology and evolutionary biology (Chomicki et al., 2019). Mutualism may be more prevalent in tropical regions than in temperate zones because the warmer and more constant climate promotes strong year-round biotic interactions (Schemske et al., 2009). Mutualism could therefore play an important role in generating the high species richness observed at low latitudes through facilitating allopatric speciation (Gómez & Verdú, 2012) or more intense coevolution with, or adaptation to, mutualistic partners (Baskett et al., 2020; Hembry et al., 2014; Weber & Agrawal, 2014). Alternatively, high species richness in the tropics could create more opportunity for species interactions to occur, and thus influence the generation and persistence of mutualism in those regions (Chomicki & Renner, 2017). Mutualisms are also expected to be more specialized in high-diversity tropical regions, promoting coexistence among many species (Schleuning et al., 2012). However, support for the hypothesis that there is more diversity and specialization in mutualisms in the tropics is mixed (Mittelbach et al., 2007; Schemske et al., 2009). Here, we investigate whether there is a latitudinal diversity gradient in the communities of rhizobia and other bacteria that live in root nodules on a legume with a large latitudinal range.

Legumes are found worldwide, live in diverse habitats, and interact with a wide range of soil microorganisms. In nutrient-poor environments, legumes depend on rhizobia to provide them with sufficient nitrogen for growth. Rhizobia fix atmospheric nitrogen in root structures called nodules in exchange for carbon and shelter from the plant. Depending on soil nutrient availability and which and how many rhizobia occur in a habitat, a legume’s dependency on symbiosis with rhizobia may vary across the globe (J. Sprent et al., 2013; J. I. Sprent & Gehlot, 2010). Tropical regions differ markedly from temperate regions in terms of soil nitrogen, phosphorus, rhizobia availability, and herbivory (Barker et al., 2022; Lambers et al., 2008), all of which could affect legume-rhizobium interactions, including host plant specificity for rhizobia.

Specificity on rhizobia varies considerably among legume species and depends on a complex exchange of signal molecules between the plant host and rhizobial partner (Hirsch & Fujishige, 2012). There is evidence that this signaling process is influenced by environmental factors, and many studies have found that tropical legumes tend to be generalists, while temperate species tend to be specialists (Andrews & Andrews, 2017; Lira et al., 2015; Perret et al., 2000). One reason tropical legumes associate with a large number of rhizobial partners may be because of the expected increase in diversity of rhizobia species in tropical soil (López-López et al., 2010). However, although there is abundant evidence that plant and animal species richness increases at lower latitudes (Jenkins et al., 2013; Mannion et al., 2014), research has failed to consistently uncover a latitudinal diversity gradient for soil bacteria (Andam et al., 2016; Fierer & Jackson, 2006). Another reason tropical and temperate legumes might differ in their number of rhizobia partners is soil nitrogen and phosphorus availability across latitude. Older and highly weathered tropical soils are expected to be phosphorus-limited compared to younger temperate soils (Lambers et al., 2008, 2009). Latitudinal patterns of nitrogen levels in soil are more complicated (Hedin et al., 2009). Some sources suggest that temperate soils are nitrogen-poor relative to tropical soils (Lambers et al., 2008, 2009; J. Sprent et al., 2013). However, tropical soil can also be somewhat nitrogen-limited due to leaching and denitrification, just not to the same extent as they are phosphorus-limited (Pajares & Bohannan, 2016). Since the legume-rhizobium mutualism is facultative and plants can survive without rhizobia when in nitrogen-rich soil, latitudinal differences in nitrogen could greatly impact legume-rhizobium mutualisms.

There are competing hypotheses for how legume specialization on rhizobia may vary with nitrogen and phosphorus levels. In phosphorus-limited conditions, adaptations for phosphorous uptake may be much more important than the legume-rhizobium mutualism (a source of nitrogen) (Werner et al., 2015). Therefore, tropical plants might nodulate less overall and only associate with a few rhizobia strains (Simms & Taylor, 2002; Tsai et al., 1993). In nitrogen-limited conditions in temperate regions, legumes may be more dependent on rhizobia since symbiosis with rhizobia is the main way temperate legumes receive nitrogen (Werner et al., 2015). Under low-nitrogen conditions, plants might associate with all available rhizobia partners to acquire any amount of nitrogen they can (Rubio Arias et al., 1999; Simms & Taylor, 2002; J. Sprent et al., 2013; Thomas et al., 2000). Thus, we might expect a gradient from generalist legumes in temperate regions to specialist legumes in tropical regions, if soil nitrogen does indeed increase towards the equator as expected.

Alternatively, under low phosphorus conditions in the tropics, plants might be less choosy of rhizobia strains and form nodules with a wide variety of rhizobia species, because choosing the “best” partner is less important and could be costly when nitrogen is not limiting (Kiers et al., 2007; Sachs & Simms, 2006). In nitrogen-poor soils in temperate regions, plants might be extremely choosy and form nodules with only the most effective nitrogen-fixing rhizobia strains (Zhang et al., 2020), especially if the investment of carbon into the mutualism is more costly under colder temperatures (Houlton et al., 2008). This would create the opposite gradient whereby temperate plants are specialists and tropical plants are generalists. We require more empirical data, especially from under-sampled tropical plants and their rhizosphere, to determine whether there are latitudinal gradients in legume specialization on rhizobia and thus the diversity of symbiotic microbes.

The microbial community found in nodules was previously thought to consist only of rhizobia that exchange signals with plants to form nodules and fix nitrogen once inside the nodule (e.g. Vincent 1970). However, recent work has uncovered non-rhizobia species (microbes that cannot independently form nodules or fix nitrogen) can enter and live inside the nodule as well (Martínez-Hidalgo & Hirsch, 2017; Selvakumar et al., 2013). The impact these non-rhizobia strains have on plant growth is extremely varied. Some studies have shown that non-rhizobia strains increase nodulation and thus plant growth (Bai et al., 2002; El-Nahrawy & Omara, 2017; Halverson & Handelsman, 1991) while others show that the non-rhizobial strains have no effect on plant performance (Bai et al., 2002). There are also varied interactions between non-rhizobia bacteria and rhizobia within nodules, ranging from mutualistic to antagonistic associations (Hansen et al., 2020). Whether or not the non-rhizobia portion of the nodule microbial community should follow a latitudinal gradient is also unclear.

We investigated how microbial diversity in *Chamaecrista nictitans* nodules changes across latitude and tested the symbiotic effectiveness of some of the strains identified in tropical and temperate regions of the range. *Chamaecrista nictitans* grows across a large latitudinal range and is found in the temperate northern region of the United States and in tropical central America. The microbial community associated with *C. nictitans* is largely uncharacterized but it has been reported to form nodules with members of the *Bradyrhizobium* genus (Koppell, 2011). We asked three main questions about the *Chamaecrista-*rhizobium system and latitude: 1) are tropical plants generalists compared to temperate plants?, 2) is the microbial community in nodules determined by microbial communities in the soil or plant host selection?, and 3) what are the effects of non-rhizobia bacteria on plant growth?

## Materials and methods

### Sample collections

We collected seed, soil, and root nodules from 33 *C. nictitans* populations across a latitudinal gradient starting in Virginia, U.S.A. (38° N) and ending in Costa Rica (8° N, Fig. 1a). Our sampling gradient covers a large part of *C. nictitan’s* native range and approaches both the northern and southern range edge (Hollenbach, 1952). We sampled 1-20 plants per population (spaced approximately 1-2 m apart to reduce our chances of sampling close relatives) in the later stages of the life cycle to collect both ripe seed and nodules with live microbes. We removed all ripe seed from each plant and stored it in dry envelopes. We dug up plant roots being careful not to damage intact nodules. Soil was shaken off the plant and collected in Falcon tubes (around 2-3 ml of soil) and the roots were rinsed or submerged in water to remove any leftover soil particles. We haphazardly removed up to 25 nodules from the roots of each plant with flame sterilized forceps and stored them in silica tubes at 4°C to preserve the microbial community. We measured and froze 2-3 g of soil (average of 2.244 g per plant) that was in close contact with the roots.

**Figure 1.**
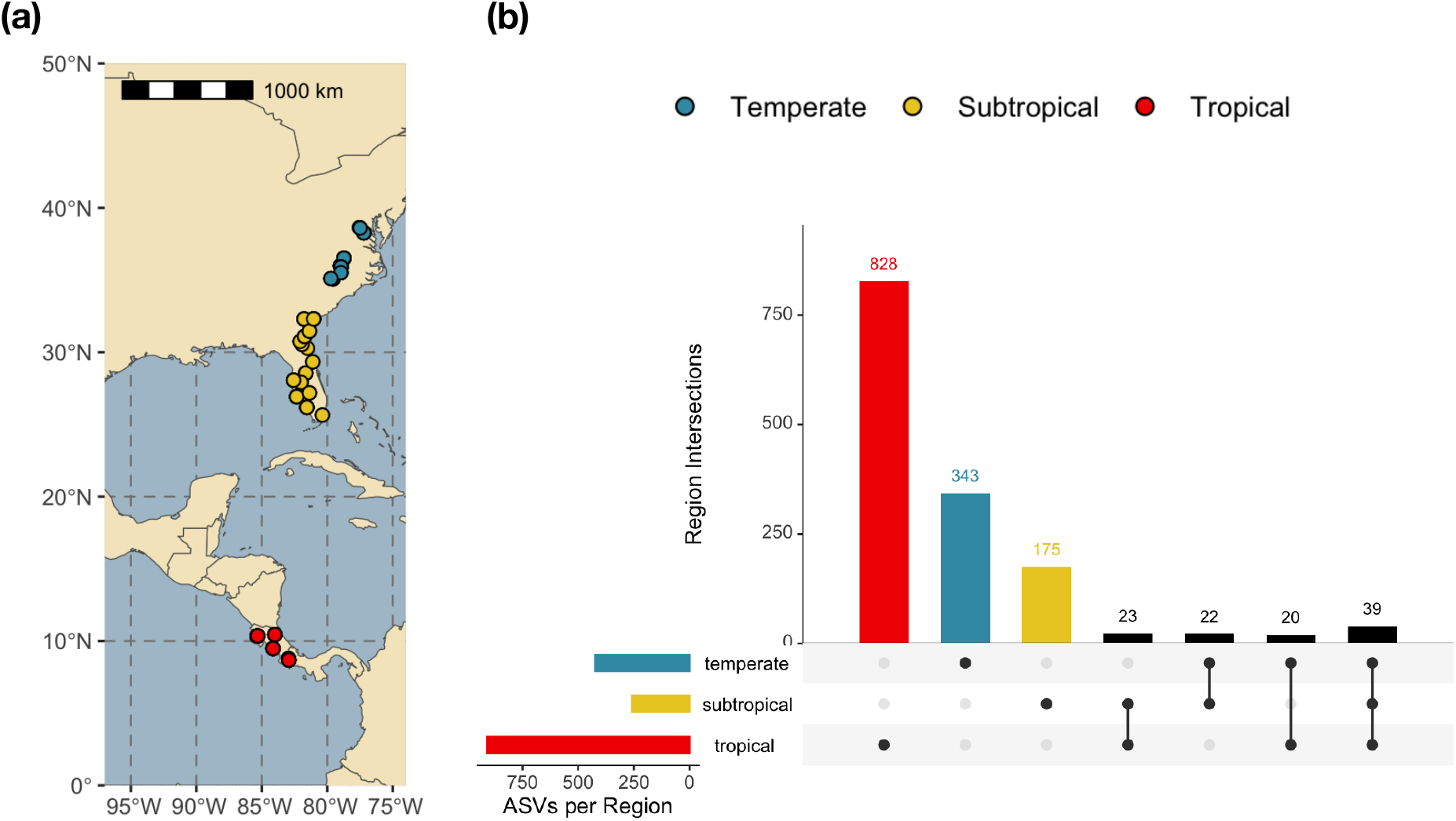
Map of sampling locations and breakdown of nodule ASVs by geographic region. a) Map of *Chamaecrista nictitans* populations sampled for seeds, nodules, and soil microbes. Each dot is a different population where 1-20 *C. nictitans* plants were sampled. Point colours show geographic region (red = tropical, yellow = subtropical, blue = temperate) which is consistent across all figures and analyses. b) UpsetR plot of unique nodule ASVs found in *Chamaecrista nictitans* plants in tropical, subtropical, and temperate regions. ASVs reported here were identified from microbial DNA extracted from field nodules (no growth phase in the lab) and sequenced at the 16S rRNA locus. Total number of ASVs in each region are reported on the left side panel. ASVs unique to each region are reported in the first three bars in the plot. ASVs shared among regions are reported in subsequent bars. Number of ASVs are indicated on the top of each bar.

### Sequencing of microbial communities

We re-hydrated all nodules per plant in a 2 ml Eppendorf tube of sterile water for one hour. The number of nodules we sampled per plant varied (1-23 nodules, *x̄* ± 1 s.e. = 7.56 ± 3.99). We sterilized the surface of the nodules by rinsing them with bleach for 20 seconds and then with 95% ethanol for 20 seconds. We washed all bleach and ethanol residue from the nodules using sterile water. To confirm that we sterilized the outside of the nodules thoroughly, we swabbed the surfaces of a subset of nodules on yeast mannitol (YM) agar plates. We incubated plates at 30°C for 3 weeks and observed no colony growth.

To extract microbial cells from nodules, we followed the methods outlined by Burghardt et al. (2018). First, we transferred the sterile nodules into a new sterile 2 ml Eppendorf tube. We added 1ml of 0.85% NaCl solution to each tube and crushed the nodules with an autoclaved pestle. We then centrifuged the solution for 10 minutes at 400 x g and pipetted off the supernatant and saved it in a separate tube. We repeated the crushing and centrifugation process in an additional 1ml of 0.85% NaCl solution and combined supernatants. We used the supernatant to create two different sample types to sequence: cultured microbes (microbes grown in the lab to high densities; see Supp. Materials and Methods) and field microbes (undifferentiated microbial cells directly from field nodules). Analysis of the cultured microbes were biased towards fast-growing taxa (Supp. Fig.1-2 and Supp. Table 1-2) and therefore we focus most of our analyses on the field samples. To extract DNA from field microbes, we used approximately 1.5 ml of the supernatant directly in the first step of the DNeasy UltraClean Microbial Kit without allowing for microbe growth in the lab. We modified the first step in the Qiagen instructions by centrifuging the supernatant for 10 minutes instead of one minute to form a sufficient pellet of microbial cells for DNA extraction. We eluted final DNA in ultrapure water. We collected DNA from 96 field samples.

To extract DNA from soil samples, we added 4 ml of sterile 0.85% NaCl solution to each tube and vortexed the solution briefly. We then centrifuged the samples at a low speed of 400 x g for 8 minutes to bring bacterial cells to the surface and soil particles to the bottom of the tube. We pipetted the supernatant to a clean 1.5 ml Eppendorf tube and centrifuged at a high speed of 15 000 x g in one-minute intervals to pellet the bacterial cells. We repeated this step until all of the supernatant was transferred to the new clean Eppendorf tube. The pellet was used as starting material in the DNeasy UltraClean Microbial Kit from Qiagen to obtain DNA. We eluted the final DNA in ultrapure water that was heated to 75°C.

We submitted 10-20 ng of DNA of field and culture samples to Genome Quebec for 16S rDNA amplification, amplicon barcoding and normalization, and library preparation for paired-end sequencing on an Illumina MiSeq PE250 platform. We chose to sequence the V3-V4 region using the primers 341f/805r because this region is hypervariable between species but highly conserved within bacterial species, and thus effective for identifying a broad range of bacteria species (Janda & Abbott, 2007). We sequenced 250 bp on two runs for a total of 253 samples (culture and field samples combined). We received a total of 24,391,826 demultiplexed reads with an average quality of 35. We sent 67 soil samples (around 10 ng of DNA each) to Genome Quebec for the same sequencing specifications. All soil samples were sequenced on one run. For soil samples, we received a total of 3,929,781 demultiplexed reads with an average quality of 36.

We identified amplicon sequence variants or ASVs (Callahan et al., 2017) by processing the reads in the program QIIME2 (Bolyen et al., 2019) using the dada2 denoise-paired method. We used default parameters for the dada2 denoise-paired method, which performs basic quality score filtering, joins paired reads, and checks for chimeras (Callahan et al., 2016). We omitted ASVs that had fewer than 20 reads total (across all samples). We identified the taxonomy of the ASVs by performing 90% similarity query searches using Blast (Bokulich et al., 2018) against the 16S 99% Greengenes database (DeSantis et al., 2006; McDonald et al., 2012). To remove potential plant contaminants from our frequency table, we excluded features that were classified as mitochondria or chloroplasts. We generated a phylogenetic tree using the align-to-tree-mafft-fasttree command with default parameters. The tree file, feature table, and taxonomy table were converted to file formats for input into R (R Development Core Team, 2020) for the rest of the data analysis. We rarefied the data with the package phyloseq (McMurdie & Holmes, 2013) to a sampling depth of 2257 reads. The final frequency table contained a total of 249 samples (93 field samples and 156 culture samples) and 2911 ASVs after all filtering procedures. We processed the soil samples using the same procedure as the field and culture samples. We combined the soil reads with their corresponding field reads (from nodules) in the QIIME2 procedure for calling ASVs. We rarefied the combined soil and field samples at a depth of 2928 reads (the size of our smallest sample) and the final frequency table contained a total of 159 samples and 20285 ASVs after filtering.

Our nodule samples (field and culture) contained a diversity of microbes including non-rhizobial species that do not nodulate or fix nitrogen. We split the data into a rhizobia community dataset and a non-rhizobia community dataset for separate analyses. To identify rhizobia ASVs, we filtered taxa for genera that are reported to be rhizobia species that make nodules and fix nitrogen in plants (de Lajudie et al., 2019, Weir 2006). We included taxa in the rhizobia dataset only if they belonged to one of the following genera: *Bradyrhizobium, Rhizobium, Mesorhizobium, Ensifer, Sinorhizobium, Cupriavidus, Burkholderia, Devosia, Microvigra, Phyllobacterium, Ochrobactrum,* and *Shinella*. Since tropical strains of rhizobia are probably not well represented in the Greengenes database and there were many ASVs in our dataset that had an unknown classification at the genus level, we also identified rhizobia ASVs by inspecting the phylogeny of the 16S rDNA sequences. Any ASVs that clustered together with known and identified rhizobia strains were also characterized as rhizobia. Alternative methods of identifying rhizobia taxa were explored (Supp. Materials and Methods) but achieved similar overall results.

### Assessment of geographic turnover in nodule communities

Because our culturing methods influenced the microbial community (see Supp Fig 1-2, Supp Table 1-2), we analyzed 1450 ASVs across 93 field samples to assess changes in nodule microbe communities across latitude within *C. nictitan’s* range. We identified the top ASVs across all samples. To assess differences in ASV frequencies in different portions of the range, we divided the data into three regions: tropical (<11° N), subtropical (>25° N and <33° N), and temperate (>35° N). We merged samples by region and calculated the frequency of each ASV in each region by dividing the number of samples the ASV was present in by the number of total samples in each region (tropical: 32, subtropical: 44, temperate: 17). We also calculated the number of unique ASVs found in each region and the total number of ASVs shared between regions.

We calculated several beta diversity indices on the field ASV data using the phyloseq package (McMurdie & Holmes, 2013). To evaluate similarities in community structure between samples we calculated Jaccard’s distance on the presence/absence ASV data and calculated Bray-Curtis distance on abundance ASV data. We also calculated unweighted UniFrac distance to take into consideration phylogenetic differences between ASVs and identify changes in unique variants across latitude. In addition, we estimated weighted UniFrac distance to identify differences in common variants. We applied a square root transformation to all distance values in order to remove negative eigen values in our principal coordinate analysis (PCoA), although raw distance values achieved similar results. We plotted the first two PCoA axes for visualization of similarities in microbial community structure across latitude. We repeated the PCoA on the rhizobia and non-rhizobia subset of the total ASV community.

### Analysis of latitudinal diversity in nodule microbe richness

To assess differences in taxa richness across latitude, we summed the raw number of unique ASVs for each sample. We performed linear or generalized linear models using the package lme4 in R (Bates et al., 2015), with the ASV counts as the response variable and latitude as the predictor variable. We used the Anova function (type III) in the car package (Fox & Weisberg, 2018) to test significance in all our models. Since the response variable was non-normal and overdispersed, we fit the taxa richness values with a quasi-poisson distribution. We ran additional models with the number of nodules included as a covariate (see Supp. Materials and Methods for details) and determined that differences in nodule number sampled among plants did not impact bacterial richness. We repeated the above analyses after removing outliers (extremely diverse samples in the tropics) from the dataset and achieved similar results, therefore we only report results from analyses performed on the full dataset. Within the rhizobia and non-rhizobia communities, we calculated ASV richness and fit linear or generalized linear models with taxa richness as the response variable and latitude as the predictor variable. In the rhizobia community analysis, we performed a linear model on the raw richness values. In the non-rhizobia community analysis, we fit a negative binomial distribution to the richness data in a glm.nb model to meet model assumptions.

To assess sequence similarity among ASVs within each geographic region (tropical, subtropical, temperate), we calculated pairwise phylogenetic distances (branch length values based on nucleotide differences in the sequence) between samples. The number of samples in each region varied with the temperate region having the fewest number of plant samples (n=17). We subsampled the subtropical and tropical groups by randomly selecting 17 samples within each region, calculated pairwise phylogenetic distances among those 17 samples, calculated the average of these pairwise distances, and repeated this process 100 times. We then plotted the average pairwise distances for each region. We also calculated phylogenetic distances separately for rhizobia taxa and non-rhizobia taxa since we expect rhizobia ASVs might be more phylogenetically similar.

### Comparison of soil and nodule microbe communities

Given the high diversity of rhizosphere microbes, we did not expect to find many exact sequence matches between the soil microbes and nodule microbes. Therefore, we relaxed the assumption that there should be a perfect match between sequences and merged all ASVs in the soil and nodule samples that were separated by a phylogenetic distance (branch length distance based on nucleotide differences) less than 0.03 to identify operational taxonomic units, or OTUs, clustered at 97% sequence similarity. We filtered the soil dataset for OTUs that were found in nodules to use for further analysis, although the full soil OTU dataset showed similar patterns in our analyses (results not reported). The filtered soil OTU dataset contained a total of 390 OTUs (from 66 soil samples) that were also found in at least one nodule sample, meaning they can gain access to nodules. We classified these OTUs into rhizobia and non-rhizobia taxa and the resulting dataset consisted of 61 rhizobia OTUs and 329 non-rhizobia OTUs.

We analyzed a total of 61 plant samples that had both a soil and corresponding nodule sample. We plotted OTU overlap across paired soil and nodule community samples to assess strain match between soil and nodule samples. We also tested for differences in the overall soil and nodule community by calculating several beta diversity metrics including Jaccard’s distance, Bray-Curtis, UniFrac, and weighted UniFrac. We performed PCoA for each of these different metrics to visualize similarities in community structure between the two sample types. We tested whether the soil community helps predict the community observed inside nodules by performing linear models on PCoA axis 1 scores from the soil samples (predictor variable) and the PCoA axis 1 scores from the corresponding nodule samples (response variable). We included population as a random factor in the models. To assess whether differences between soil and nodule communities varied across latitude, we fit linear models with raw UniFrac or weighted UniFrac distance (calculated between soil and nodule samples) as the response variable and latitude as the predictor variable.

### Analysis of differential diversity and abundance of soil microbes across latitude

We performed linear models on the soil sample data with OTU richness as the response variable and latitude as the predictor variable. We included only OTUs that can occupy nodules, rather than using the entire soil community including strains that do not interact with plants. We performed linear models on the full community, rhizobia only community, and non-rhizobia only community. Raw OTU richness counts were used in all models since they were normally distributed. To identify differentially abundant rhizobia and non-rhizobia genera between tropical, subtropical, and temperate soils, we performed an analysis of compositions of microbiomes with bias correction (ANCOMBC, Lin & Peddada, 2020). We conducted ANCOMBC models on raw (non-rarefied) soil data filtered for OTUs that can occupy nodules.

### Co-inoculation experiments of rhizobia and non-rhizobia strains

We identified many non-rhizobia species within *C. nictitans* nodules (Supp. Fig. 2). Therefore, we tested the effects of some of these non-rhizobia species on plant growth by conducting growth chamber and greenhouse experiments at the University of Toronto in November 2019. Specifically, we co-inoculated *C. nictitans* plants with non-rhizobia and rhizobia strains isolated from field nodules. We identified and isolated the rhizobium species *Bradyrhizobium elkanii* and two non-rhizobia species *Pseudomonas korensis* and *Variovorax paradoxus* from a single temperate plant (Supp. Materials and Methods). From the opposite end of the *C. nictitans’* range, we isolated one rhizobium species, *Bradyrhizobium yuanmingense,* and one non-rhizobia species, *Bacillus cereus,* from a single tropical plant (Supp. Materials and Methods). Because traditional culture-based methods for isolating bacteria often fail to capture the most abundant microbes in field samples (O’Brien et al., 2020), the prevalence of the non-rhizobia strains used in our experiments was low across the nodule (field) samples in our ASV dataset. We found a total abundance of 113 reads of the *V. paradoxus* species, a total of 18 reads of the *B. cereus* species, and a total of 341 reads of the genus *Pseudomonas* (we were unable to classify *Pseudomonas* ASVs to the species level to identify *Pseudomonas koreensis*). *Bacillus cereus* was the most common species among the cultured microbes (Supp. Table 1). We conducted a greenhouse experiment testing the effects of the temperate derived strains on field-collected seeds from the source temperate population and a growth chamber experiment to test the effects of the tropical strains on field-collected seeds from the source tropical population (see Supp. Materials and Methods for greenhouse set up).

We planted a total of 123 plants in a full 2x2 factorial design where we manipulated the presence of *Bradyrhizobium yuanmingense* and *Bacillus cereus* in the tropical strain experiment. Each plant was assigned one of four treatments: 1 ml of rhizobia *B. yuanmingense* + 1 ml control, 1 ml of non-rhizobia *B. cereus* + 1 ml control, 1 ml rhizobia + 1 ml non-rhizobia, or 2 ml of control media (clean YM liquid media). We prepared strains by growing the microbial samples in liquid YM cultures. Magenta Boxes containing single plant individuals were randomized across the growth chamber floor. Seeds used in the experiment were collected from five separate maternal plants from the tropical source population. A total of 28 plants across all four treatments died early in the experiment before microbe cultures were added and were thus removed from analysis.

A total of 270 seeds were planted in a randomized complete block design on a single bench in a greenhouse for the temperate experiment. All maternal plant families (12 separate maternal plants from the source temperate population) and experimental treatments were represented in each block. We manipulated the presence of *B. elkanii, P. korensis,* and *V. paradoxus* in a 3 x 3 factorial design where each plant was assigned one of eight treatments: 3 ml control, 1 ml *B. elkanii +* 2 ml control, 1 ml *P. korensis +* 2 ml control, 1 ml *V. paradoxus* + 2 ml control, 1 ml *B. elkanii* + 1 ml *P. korensis* + 1 ml control, 1 ml *B. elkanii + V. paradoxus* + 1 ml control, 1 ml *P. korensis +* 1 ml *V. paradoxus* + 1 ml control, and finally 1 ml *B. elkanii* + 1 ml *V. paradoxus* + 1 ml *P. korensis.* We cultured the three strains using the same methods described for the tropical experiment (Supp. Materials and Methods). A total of 54 plants died early in the experiment before treatments were applied and were removed from the data analysis.

In both experiments, plants were harvested after approximately four months of growth for measurements of aboveground biomass, belowground biomass, nodule number, and nodule weight. All nodules on fresh plant roots (stored at 4°C) were counted under a microscope within a week of harvest. We randomly collected up to five nodules from the roots and dried them in tubes with silica gel for 2 weeks. We measured the dried nodules and then divided the values by the number of nodules measured to get an average weight per nodule. We then separated roots from shoots and dried the plant material at 60°C for three days before measuring belowground and aboveground weight in mg. Leaf counts and mortality were scored on the plants at multiple points throughout the experiment.

We analyzed aboveground biomass and nodule number using linear and generalized linear mixed models for both experiments and nodule weight for the tropical experiment. We included maternal plant family as a random effect in the analyses of data from both experiments and block as a random effect in the temperate analysis. We log_10_ transformed aboveground biomass to meet assumptions of normality. We fit strain presence as the predictor variable and included an interaction between *B. cereus* and rhizobia in the tropical experiment and two- and three-way interactions between rhizobia, *P. korensis,* and *V. paradoxus* in the temperate analysis. We fit the nodule number response variable to a poisson distribution with link set to “sqrt” and included plant identity as a random factor to account for overdispersion and meet model assumptions in the tropical experiment. To determine if the addition of *B. cereus* increased variance in nodule number, we performed a Bartlett’s test comparing rhizobia treatments with and without *B. cereus*. Nodule weight in the tropical experiment was only measured on plants that formed nodules, which included only plants in the rhizobia treatments. Therefore, we ran a linear mixed model testing the impact of *B. cereus* on average nodule weight for only the experimental treatments with rhizobia. Nodule number data in the temperate experiment was zero-inflated and overdispersed, however zero-inflated models with a poisson distribution or negative binomial were over-parameterized and did not perform well. Therefore, remaining analyses on the nodule number parameter were performed only on treatments that contained plants with nodules (rhizobia treatments). We fit the nodule number response variable to a poisson distribution with link set to “sqrt” and included plant identity as a random factor in a generalized linear mixed model. We tested for an interaction between *P. korensis* and *V. paradoxus*. We also compared the variance in nodule number between the rhizobia treatments with and without the *P. korensis* and *V. paradoxus* strains using a Levene’s test since the data were not normally distributed.

## Results

### High turnover in nodule microbe communities across latitude

Although *C. nictitans* plants are mainly nodulated by microbes in the genus *Bradyrhizobium* across the species’ range (Supp. Table 3), there is turnover with latitude in which specific *Bradyrhizobium* ASV strains are present in *C. nictitans* nodules. Different *Bradyrhizobium* ASVs were found in all three regions at different frequencies, so no single ASV was ubiquitous across populations. The most common ASV (*Bradyrhizobium*, species unidentified) across the field samples was present in 51.62% of the samples. The subtropical and temperate regions shared the same top five most common ASVs (Supp. Table 3). The most prevalent ASV in the tropical region was found in 84.4% of tropical samples, but was absent from all temperate samples, and found in only 29.5% of subtropical samples. We also saw that the tropical region contained the highest number of unique ASVs overall (Fig. 1b). The most common ASV in the temperate/subtropical zones was found in 76.5% of temperate plants, 77.3% of subtropical plants, and only in 3.1% of tropical plants (Supp. Table 3). Only a small number of ASVs (39) were shared among the different geographic regions (Fig. 1b). PCoA plots based on Bray-Curtis distance (Fig. 2a) and Jaccard’s distance (Supp. Fig. 3a) also show a strong separation of tropical and temperate plant samples on axis 1 in the full community, but the breakdown of rhizobia and non-rhizobia ASVs (Fig. 2b-c, Supp. Fig. 3b-c) show that this is largely due to differences in rhizobia strains across latitude. When we look at dissimilarity metrics that take into account phylogenetic similarity in the microbial community, there is less clustering of the samples in PCoA space. Therefore, although microbe strain diversity was high and varied across the range, *C. nictitans* plants appear to associate with phylogenetically similar nodule communities in the overall community (Fig. 2d, Supp. Fig. 4a). Nonetheless, tropical plant samples have rhizobia communities that are clearly separated from those of temperate and sub-tropical plants along axis 2 for both UniFrac and weighted UniFrac analyses (Fig. 2e, Supp. Fig. 4b). Non-rhizobia communities show no obvious clustering of populations when phylogenetic similarity was taken into account (Fig. 2f, Supp. Fig. 4c).

**Figure 2.**
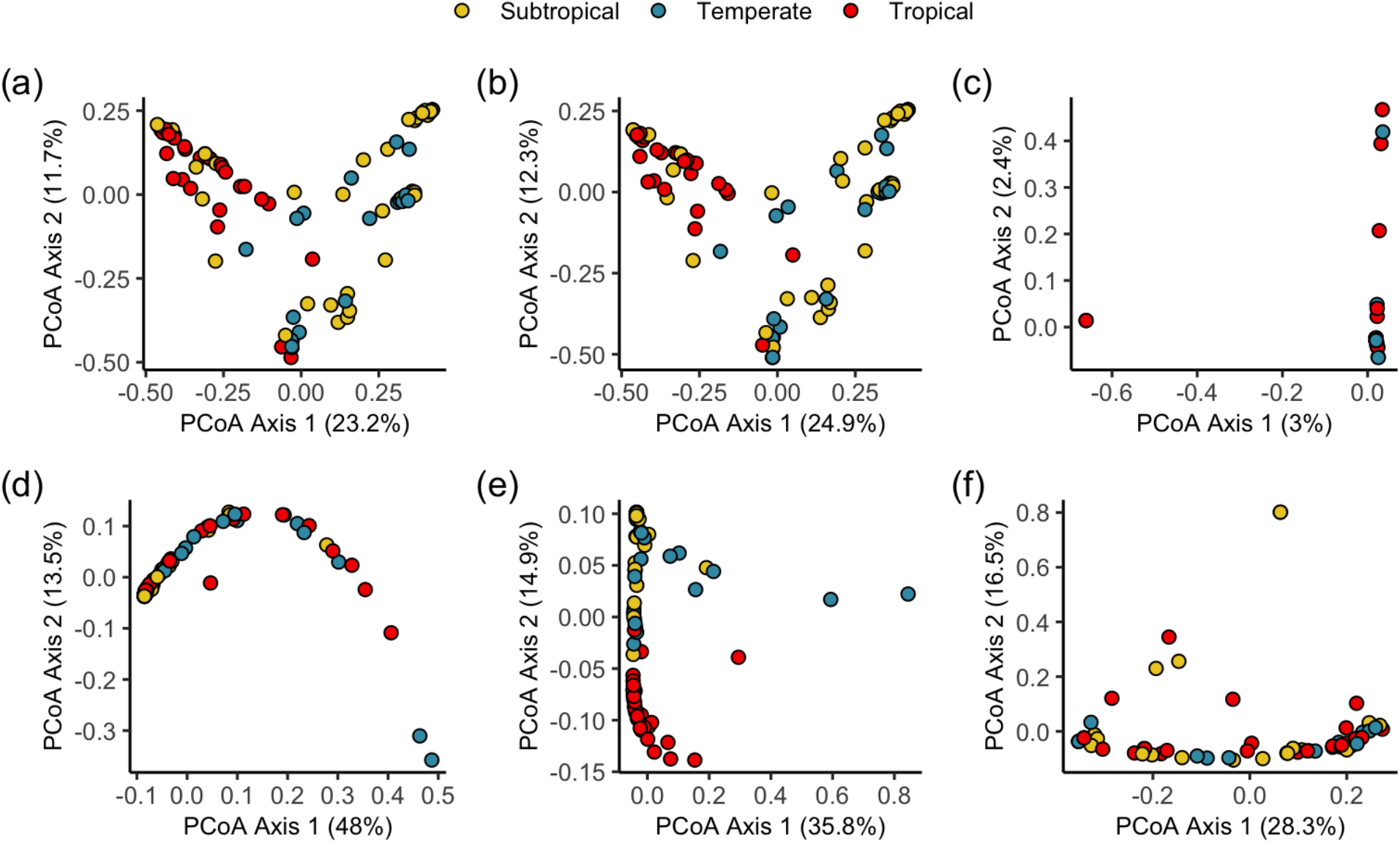
Principal Coordinate Analysis (PCoA) of nodule microbial community structure. PCoAs were plotted using Bray-Curtis (a-d) and weighted UniFrac (d-f) dissimilarity in nodule communities. Each dot represents a single *Chamaecrista nictitans* plant where nodules were pooled and the microbe community was extracted and sequenced directly (labeled “field” in further analyses) without a growth phase in the lab. Panels (a) and (d) represent PCoA performed on the full ASV community. The middle panels (b and e) represent rhizobia only ASVs and the last column of plots (c and f) are non-rhizobia ASVs.

### Latitudinal diversity gradient in nodule communityh

We observed high ASV richness at lower latitudes when we considered the entire nodule community and the non-rhizobia portion of the community (Fig. 3 top panel, Table 1). In contrast, plants were associated with similar numbers of rhizobia ASVs across the range. Pairwise phylogenetic distance averages were low in the subtropical region compared to temperate and tropical regions when we considered the whole nodule community (Fig. 4). There was greater diversity among the non-rhizobia microbe strains across all three regions compared to the rhizobia strains (Fig. 4). Rhizobia ASVs were more phylogenetically similar in the subtropical region compared to tropical and temperate regions. Non-rhizobia communities displayed similar (but high) pairwise distance averages in all geographic regions, with the exception of having more variance in the subtropical zone.

**Figure 3.**
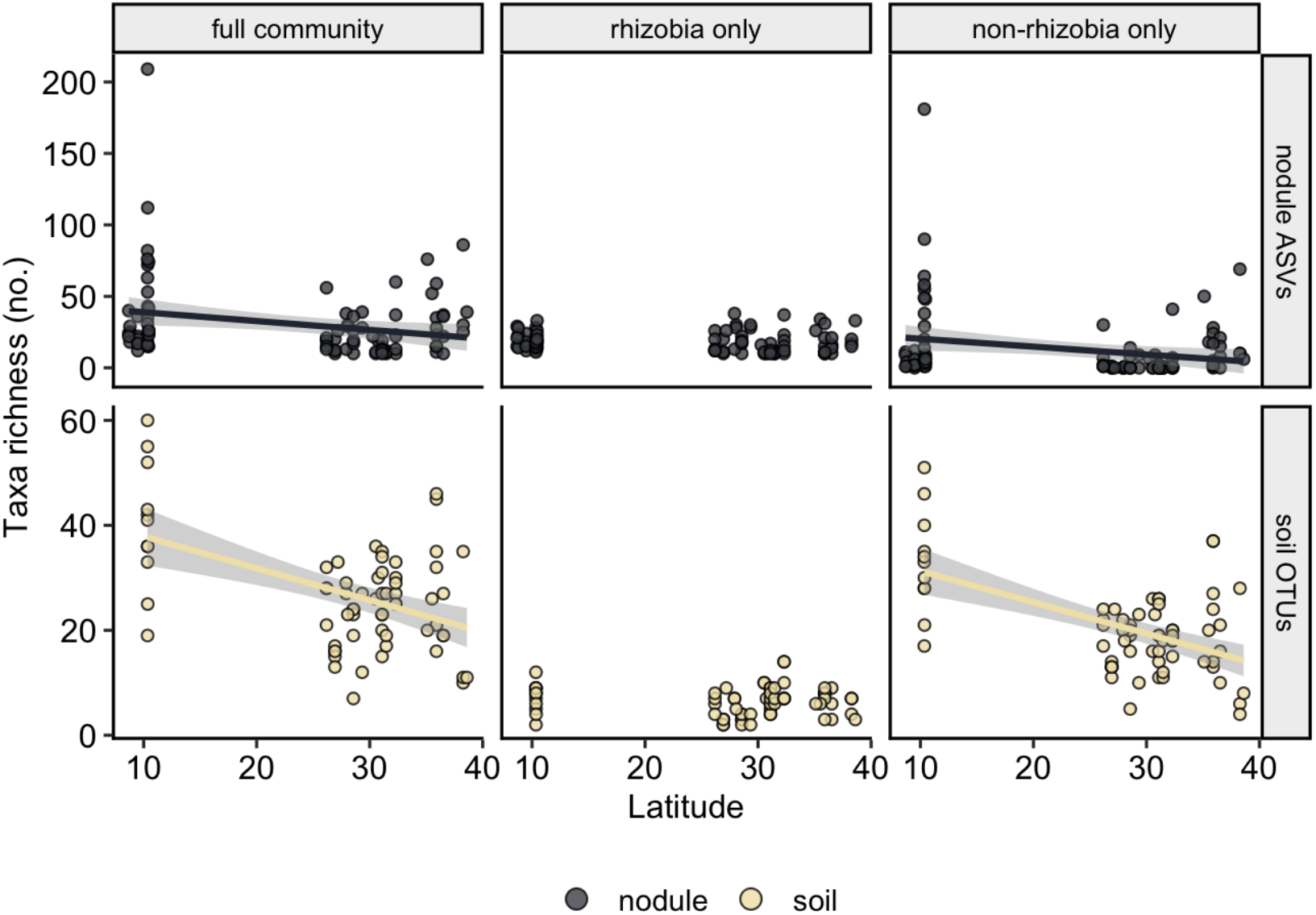
Relationship between latitude and the total number of unique rhizobia and non-rhizobia taxa. Taxa richness is reported in plant nodules of *Chamaecrista nictitans* plants (dark grey points) and the soil surrounding plant roots (light yellow points). Unique exact sequences or ASVs (nodule samples) and sequences merged at 97% similarity or OTUs (soil samples) are reported as taxa richness values. Relationships are estimated for taxa in the full community, rhizobia portion of the community, and non-rhizobia portion of the community. Taxa richness reported for the soil samples were calculated from taxa that are shared between soil and nodule samples (rather than the whole soil community). Relationships that are significant at p<0.05 have the regression line plotted on the graph.

**Figure 4.**
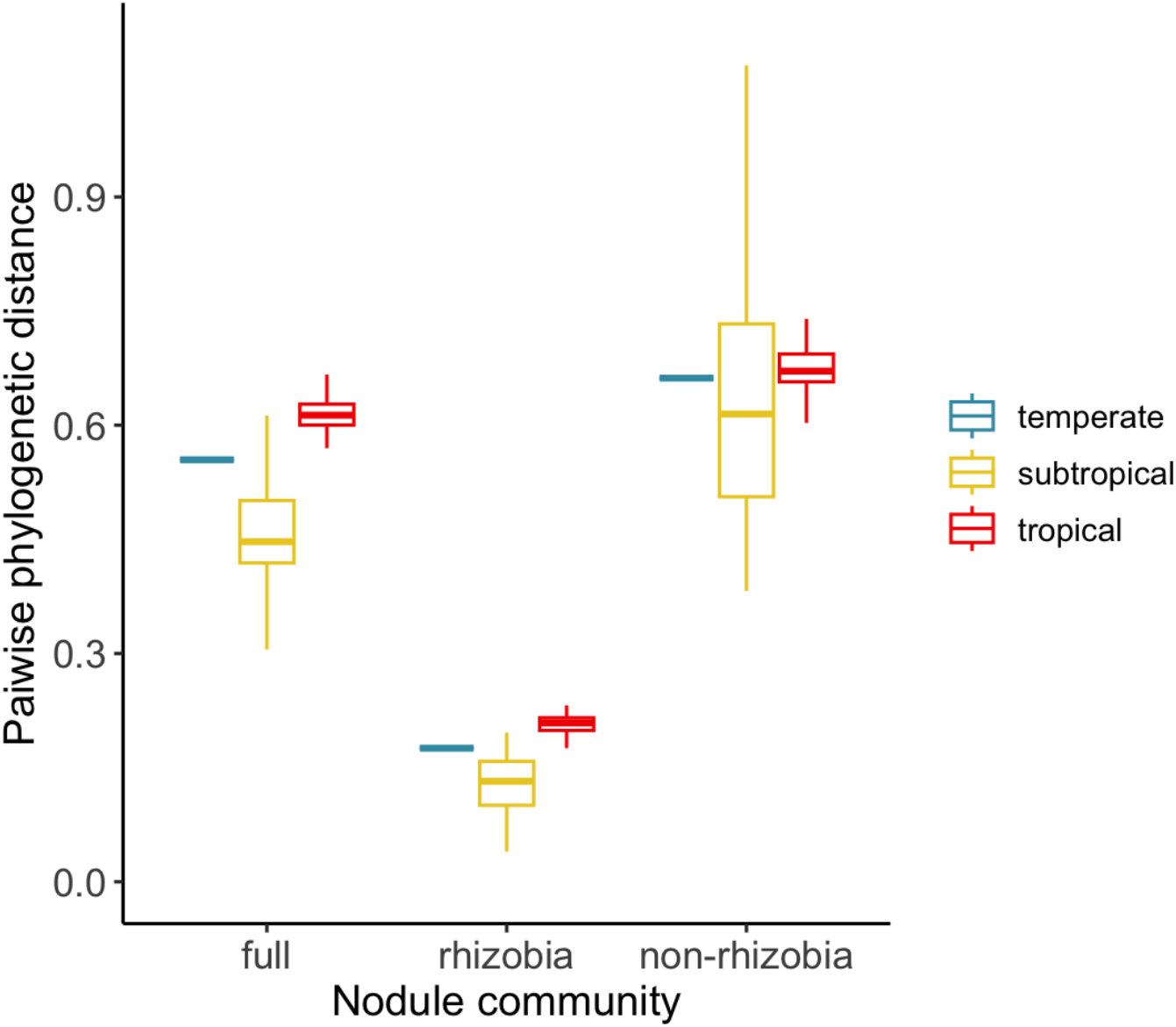
Average pairwise phylogenetic distances between nodule ASVs from temperate, subtropical, and tropical regions. Pairwise distances represent the similarity in the 16S sequence of two ASVs. Data plotted is the average of all pairwise distances for that geographic region. Phylogenetic distances shown for pairs of ASVs from the full community, rhizobia only ASVs, and non-rhizobia only ASVs. Subtropical and tropical regions were subsampled 100 times to 17 samples to match the sample size of the temperate region. Outliers (1.5 x inter quartile range) have been removed from plot for better visualization of data.

**Table 1.** Results of linear and generalized linear models testing for a latitudinal diversity gradient in the nodule and soil microbe communities. Results reported for the ASV nodule and OTU soil datasets are broken down into the full, rhizobia-only, and non-rhizobia-only communities. OTUs used in the soil analyses are taxa that are shared between soil and nodule samples. Bolded results are significant at p < 0.05. F/Wald χ^2^ values and p values are output from Anova type III tests. Wald χ^2^ statistics reported for the full and non-rhizobia community in the nodule dataset, all others reported are F values. The response variable for all tests is the number of unique ASVs.

| | estimate | s.d. | F/Wald $\chi^2$ | d.f. | p value |
| --- | --- | --- | --- | --- | --- |
| *Nodule ASVs, n = 93* |  |  | *F/Wald $\chi^2$* |  |  |
| **Full community** | **-0.0415** | **0.0153** | **7.5586** | **1** | **0.0060** |
| Rhizobia | -0.0666 | 0.0692 | 0.9258 | 1 | 0.3385 |
| **Non-rhizobia** | **-0.0337** | **0.0171** | **5.1226** | **1** | **0.0236** |
| *Soil OTUs, n = 66* |  |  | *F* |  |  |
| **Full community** | **-0.6081** | **0.1373** | **19.613** | **1** | **<0.0001** |
| Rhizobia | -0.0120 | 0.0407 | 0.0877 | 1 | 0.7681 |
| **Non-rhizobia** | **-0.5960** | **0.1111** | **28.762** | **1** | **<0.0001** |

### Latitudinal patterns of microbe diversity in soil

Only 0.19% of soil ASVs were also found in nodule samples (Fig. 5a). When we clustered ASVs by 97% similarity, we found that 4.55% of soil taxa were also found in nodules. When soil ASVs were subsetted to ones that can enter nodules, we observed a significant latitudinal gradient in the whole soil community and non-rhizobia soil community (Fig 3 bottom panel). In contrast, the rhizobia-only richness showed a weak non-significant relationship with latitude, similar to the results of the nodule community. However, this result does not mean that the latitudinal gradient in non-rhizobia microbes living inside nodules is because plants are merely sampling their soil community. The soil community did not predict the nodule community in any of our analyses.

**Figure 5.**
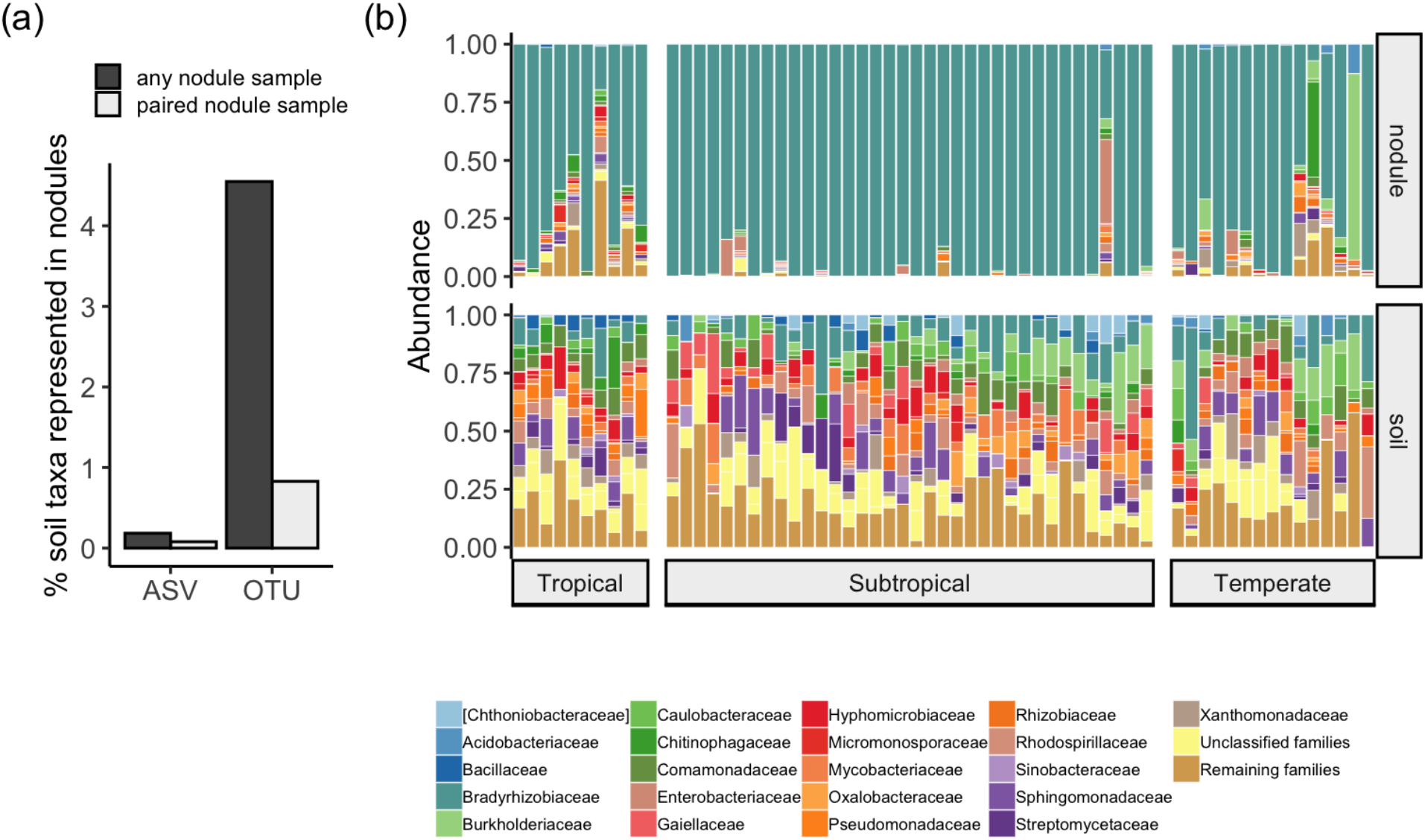
Comparison of soil and nodule microbial communities. a) Proportion of exact soil sequences (ASVs) or clustered sequences by 97% similarity (OTUs) that were also found in a matching nodule sample (light grey) or any nodule sample (dark grey). b) Differences between the microbial community collected from nodules (top) and the surrounding soil (bottom) of *Chamaecrista nictitans* plant roots. A total of 61 plants are plotted on the x-axis where each bar represents an individual plant and the bacterial community associated with that plant. ASVs were merged by the taxonomic family. Relative proportions of each microbe family are plotted for either the nodule community or soil community. The most common bacteria families that make up greater than 15% of total sequences are plotted, all other families were grouped into one category called “remaining families” (light brown). ASVs that are unidentified are represented by “unclassified families” (yellow).

Nodule samples were largely dominated by nodulating OTUs belonging to the Bradyrhizobiaceae family, while soil samples taken from surrounding the plant (and nodules) were highly diverse, containing rhizobia and non-rhizobia strains and a number of unidentified ASVs (Fig. 5b). Soil and nodule communities showed clear differences in PCoA plots for all beta diversity metrics we tested (Supp. Fig. 5a-d), but particularly with metrics that accounted for phylogenetic distance (Fig. 6a). The soil community was not a significant factor describing nodule community composition for any of the distance metrics we used in our analysis (Fig. 6b, Supp. Table 4). Differences between the soil and nodule communities were similar across the range for all beta distance metrics except unifrac distance (Fig. 7a-b, Supp. Table 5), suggesting that tropical communities hosted by plants differ more in rare variants compared to temperate microbial communities.

**Figure 6.**
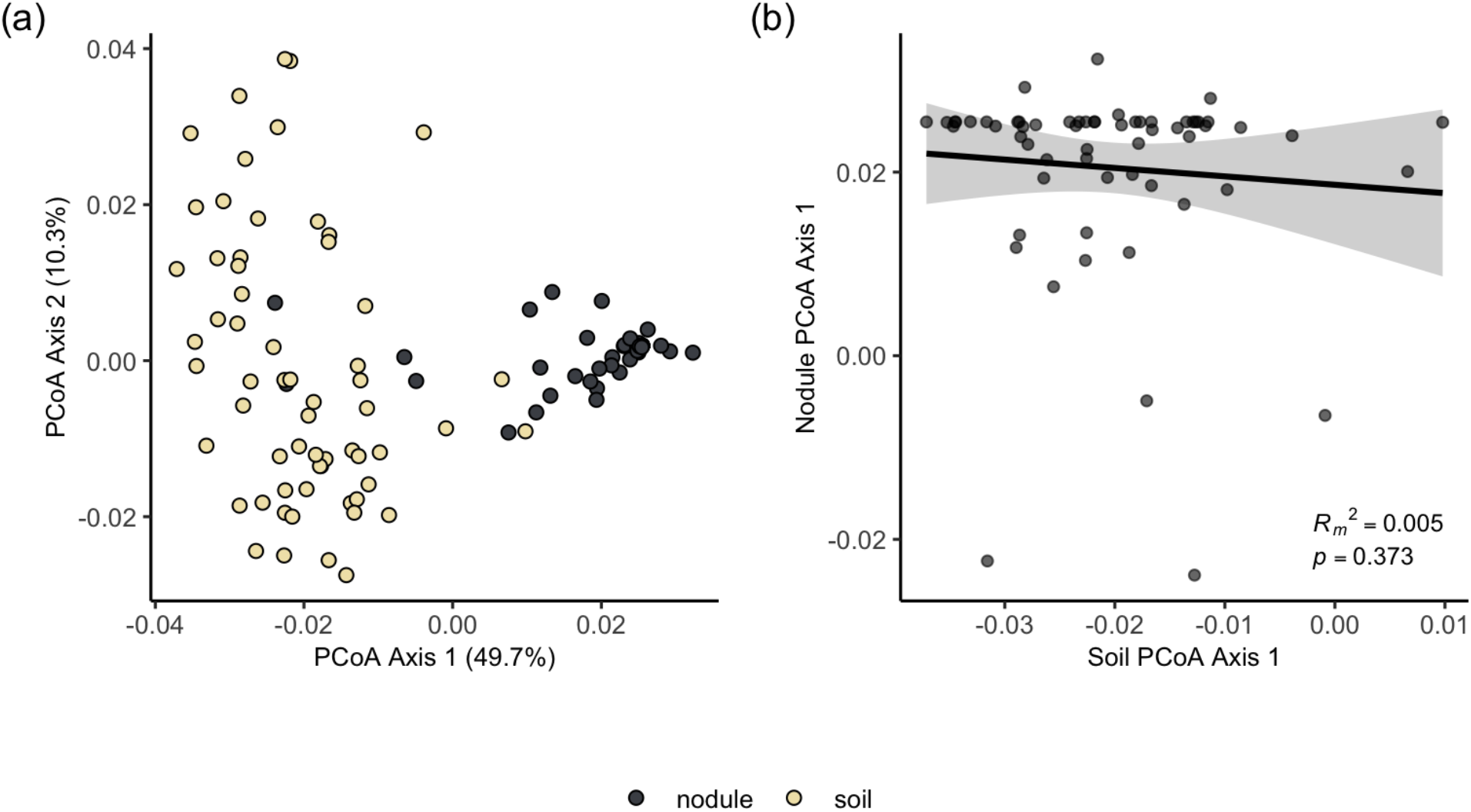
Community composition of paired soil and nodule samples. a) PCoA performed on weighted UniFrac distances calculated from nodule-isolated microbes (dark grey) and soil microbes (light yellow). Soil microbes represent only those OTUs (sequences clustered at 97% similarity) that are found in nodules. b) Relationship between soil PCoA axis 1 scores and nodule PCoA axis 1 scores calculated from weighted UniFrac distance values. Marginal R^2^ value and p value reported on the plot are results of generalized linear models testing the impact of soil PCoA axis scores on nodule PCoA axis scores with population modeled as a random factor. Each point on the plot represents one plant sample and the comparison between the PCoA scores on the surrounding soil and matching nodule.

**Figure 7.**
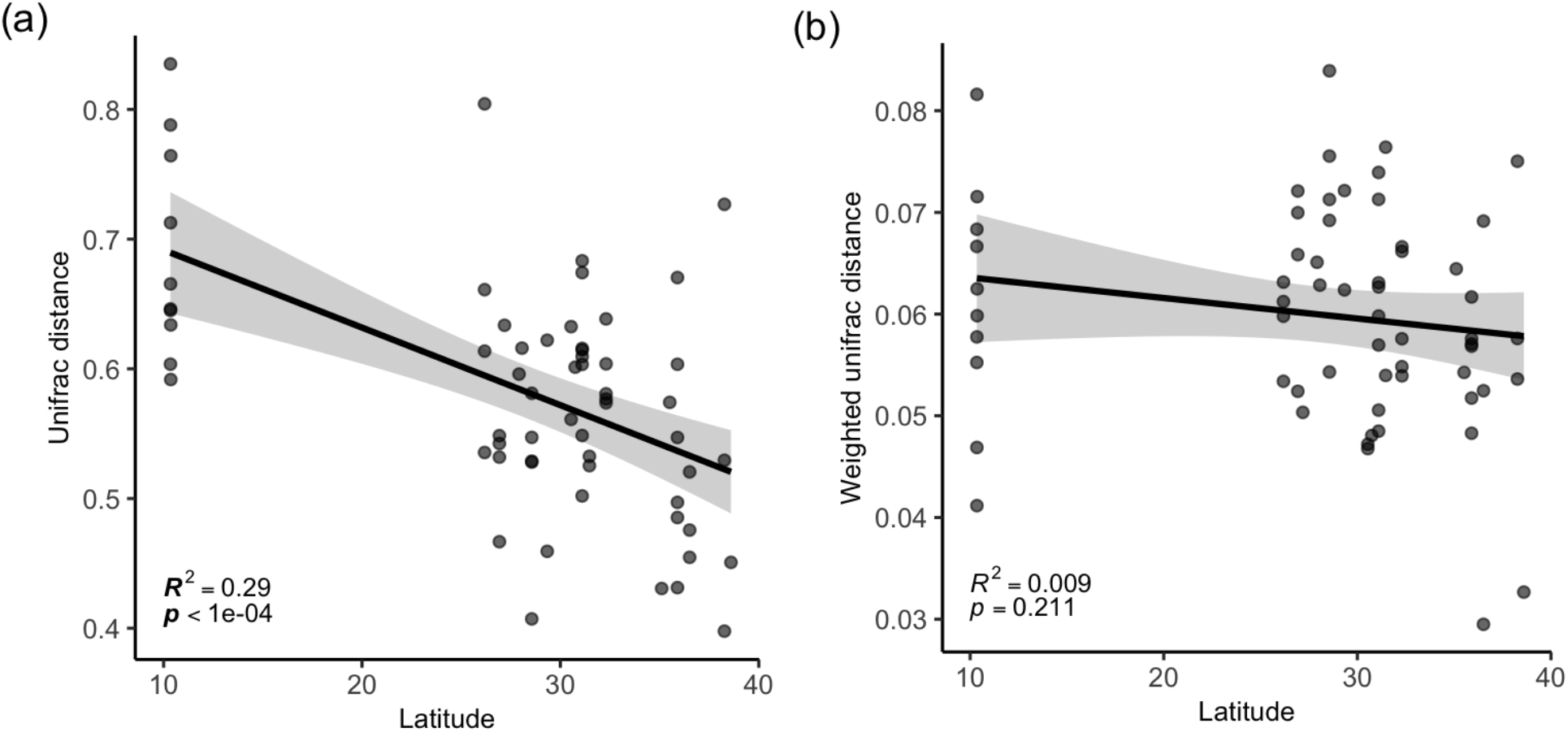
Relationship between latitude and community similarity between soil and nodule microbe communities. Each point on the plot is a plant sample where either a) UniFrac or b) weighted UniFrac distance was calculated between matching nodule and soil microbial community. R^2^ values and p values reported on the plot are results from linear models testing latitude as a predictor of community similarity. Significant relationships are indicated by p values in bold text.

Of the 35 known microbe families tested, 20 were significantly differentially abundant across the three geographic regions. Only one of these 20 families was a family known to contain nodulating rhizobia strains (Burkholdericeae). Bradyrhizobiaceae, the most common family that we found to nodulate *C. nictitans*, did not differ in read abundance across latitude. The nodulating family Burkholdericeae was fairly low in abundance in the tropics compared to the other regions (Supp. Fig. 6). The tropical region contained an even combination of non-rhizobia families that were highest (9) and lowest (9) in abundance across the range (Supp. Fig. 6).

### Non-rhizobia effects on plant growth

Overall, the addition of non-rhizobia microbes had very little impact on plant growth and nodule formation. We found no significant effect of non-rhizobia inoculations on aboveground biomass in either the tropical (Fig. 8, Table 2) or temperate experiment (Fig. 8, Table 3). There was no significant difference in variance in nodule number between the rhizobia treated plants with and without *B. cereus* (Bartlett’s K^2^=0.1139, p=0.7357). The addition of *B. cereus* also had no effect on nodule weight. The interaction between rhizobia, *P. korensis,* and *V. paradoxus* in the temperate experiment had a non-significant effect on all measurements (Table 3). The interaction between *P. korensis* and *V. paradoxus* was also non-significant in the Levene’s test comparing variances in nodule number (F=0.5045, p=0.6802).

**Figure 8.**
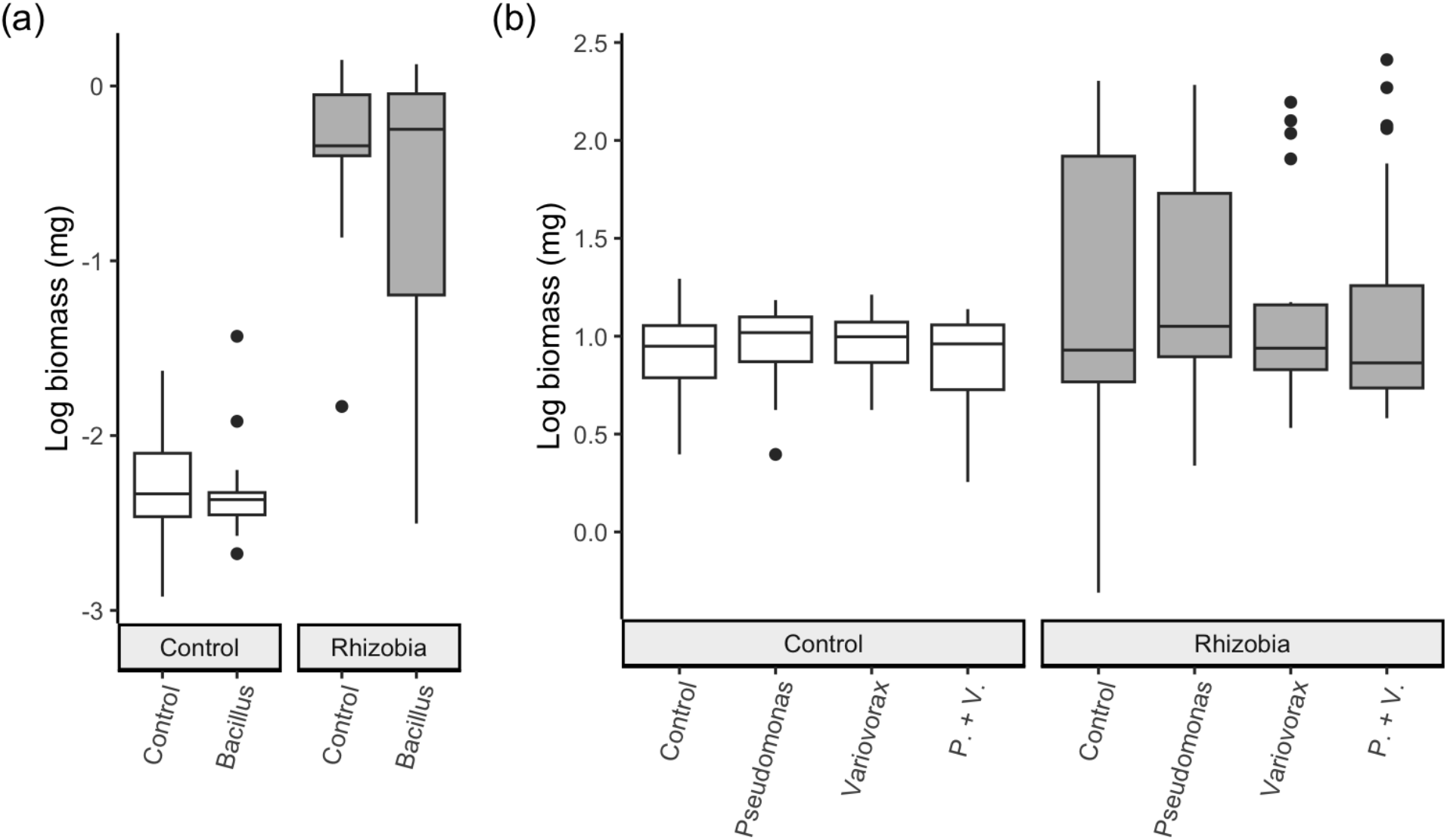
Results from the tropical (a) and temperate (b) growth chamber and greenhouse experiments of co-inoculations of rhizobia and non-rhizobia strains. The impact of the tropical *Bacillus cereus* non-rhizobia strain on aboveground biomass (log_10_ transformed) is shown in (a) and the impact of different non-rhizobia strains on aboveground biomass (log_10_ transformed) shown in (b). Treatments in dark grey are treatment plants with rhizobia inoculations.

**Table 2.** Results of mixed effect linear and generalized linear models for testing the effects of rhizobia and *Bacillus cereus* on tropical *Chamaecrista nictians* plants. Wald χ^2^ values and p values are the results of type III Anova tests. Bolded results are significant at p < 0.05. Sample size for the nodule weight data was n=35. Sample size for the remainder of the tests was n=64.

| | treatment | Wald $\chi^2$ | d.f. | p value |
| --- | --- | --- | --- | --- |
| Biomass | **Rhizobia** | **186.1172** | **1** | **<0.0001** |
|  | Bacillus | 1.2707 | 1 | 0.2596 |
|  | Rhizobia*Bacillus | 1.0205 | 1 | 0.3124 |
| Nodule no. | **Rhizobia** | **184.0669** | **1** | **<0.0001** |
|  | Bacillus | 0.5579 | 1 | 0.4551 |
|  | Rhizobia*Bacillus | 2.2612 | 1 | 0.2327 |
| Nodule weight | Bacillus | 0.0477 | 1 | 0.8271 |

**Table 3.** Results of mixed effect linear and generalized linear models for testing the effects of non-rhizobia strains on temperate *Chamaecrista nictitans* plants. Wald χ^2^ values and p values are the results of type III Anova tests. Bolded results are significant at p < 0.05. Sample size for biomass data was n=164. Sample size for the nodule number data was n=86.

| | treatment | Wald $\chi^2$ | d.f. | p value |
| --- | --- | --- | --- | --- |
| Biomass | **Rhizobia** | **9.4354** | **1** | **0.0021** |
|  | Pseudomonas | 0.1119 | 1 | 0.7380 |
|  | Variovorax | 0.3810 | 1 | 0.5371 |
|  | Rhizobia*Pseudomonas | 0.1490 | 1 | 0.7000 |
|  | Rhizobia*Variovorax | 0.0480 | 1 | 0.8266 |
|  | Pseudomonas*Variovorax | 0.0042 | 1 | 0.9486 |
|  | Rhizobia*Pseudomonas*Variovorax | 0.1066 | 1 | 0.7441 |
| Nodule no. | Pseudomonas | 0.4502 | 1 | 0.5023 |
|  | Variovorax | 0.1506 | 1 | 0.6980 |
|  | Pseudomonas*Variovorax | 0.3855 | 1 | 0.5347 |

## Discussion

We combined field sampling, microbial community analysis, and greenhouse experiments to determine whether there is a latitudinal diversity gradient within nodule microbes and their effects on plant performance. We uncovered three main results. First, we found a latitudinal gradient in bacterial richness in nodules, but only in the non-rhizobia portion of the nodule community. Second, microbial communities in the surrounding soil did not predict nodule communities. Finally, we tested a few non-rhizobia strains isolated from tropical and temperate field nodules in greenhouse experiments and found that they are most likely commensals.

### A latitudinal diversity gradient in non-rhizobia bacteria in Chamaecrista nictitans nodules

Individual *C. nictitans* plants hosted a high diversity of microbes at the strain level (ASVs) across the species’ native range. Although rhizobia strains were present in higher abundances (Fig. 5b), non-rhizobia strains were found in most samples and this portion of the community was incredibly taxonomically diverse. *Chamaecrista nictitans* has a large geographic range and legumes with large ranges tend to be generalists in their associations with rhizobia partners (Harrison et al., 2018). *Chamaecrista nictitans* seems to be a generalist in terms of selecting rhizobia species across its range in North and Central America as plants generally associated with a similar number of nodulating *Bradyrhizobium* and other rhizobia strains regardless of location. Although we did not find any relationship between latitude and the total number of unique rhizobia strains (Fig. 3), we did see high turnover in the rhizobia community across latitude (Fig. 2). There was more sequence variation among tropical ASVs compared to subtropical and temperate ASVs, even in rhizobia strains, although the non-rhizobia community was more phylogenetically diverse (Fig. 4). Rhizobia have a free-living stage in soil during which time they are not associated with a legume host. The consistent and benign year-round soil conditions of the tropics may promote microbial growth and thus foster more rapid divergence among rhizobia when in soil (Dennis et al., 2019). In addition, adaptation to a highly diverse leaf litter and other host plants could generate diverse microbial communities (Bradford et al., 2017) and higher temperatures could increase molecular rates of evolution in microbes (Wright et al., 2006). Tropical rhizobia may have also had more time to diverge considering that temperate zones would have experienced periods of continental glaciation (Brown, 2014).

We did uncover a latitudinal diversity gradient in the non-rhizobia portion of the nodule community in *C. nictitans* plants (Fig. 3). It is unclear how non-rhizobia species enter a nodule given that they generally lack *nod* genes and are unable to start a signaling exchange with plants. If plants are able to select which non-rhizobia strains enter their nodules, then host selection on the non-rhizobia community may vary across latitude. Under high-nitrogen conditions in the tropics, plants may allow more non-rhizobia strains to enter nodules because these plants do not require fixed nitrogen from rhizobia. Another possibility is that non-rhizobia cells hitch-hike into a nodule alongside a rhizobium and the plant host has no control over selecting which non-rhizobia strains achieve entry. In this case, nodule communities associated with *C. nictitans* plants could be largely determined by the surrounding soil community. Plants may host more non-rhizobia strains in the tropics simply because tropical soil contains a higher diversity or density of non-rhizobia microbes.

### Mismatch between soil and nodule microbial communities

Soil analyses showed that microbial communities found in nodules are not simply a reflection of microbes found in the soil. Although we found a significant latitudinal diversity gradient in the non-rhizobia portion of the soil community, matching the diversity patterns observed in the nodule community (Fig. 3), soil and nodule microbial microbes showed striking differences in community structure (Fig. 6) especially in the tropics (Fig. 7). The latitudinal diversity gradients in the soil and nodules may be driven by similar abiotic factors, but hosts are clearly selective on non-rhizobia as the soil community did not effectively predict the nodule community. It was surprising that so few nodule ASVs were found in the soil environment (Fig. 5a). Low abundant microbial strains in the soil (so low to the point of being undetectable by 16S rDNA sequencing) may grow to high abundances once in a more favourable environment such as inside a nodule, explaining the high mismatch between soil and nodule samples. Tropical conditions are also expected to be more suitable for microbes to grow to high abundances. If non-rhizobia strains grow to larger population sizes in the tropics, it may be easier for them to gain access to root nodules. We found an even mix of non-rhizobia families that are high and low in abundance in tropical soil compared to subtropical and temperate regions. The highly abundant families were not necessarily more represented in tropical nodules. Although we still require more research into the mechanisms of non-rhizobia bacteria access to nodules, our results suggest that host selection is important in structuring the non-rhizobia community in nodules.

### Non-rhizobia strains are commensal under controlled conditions

Since the latitudinal patterns of diversity largely occurred in the non-rhizobia portion of the nodule community, we aimed to test the effect these strains have on plant fitness. The tropical *B. cereus* and temperate *P. koreensis* and *V. paraxoxus* strains appear to be commensals based on our results. Plants inoculated with non-rhizobia strains (and no rhizobia) showed no difference in biomass compared to control plants (Fig. 8) even though previous research suggests that *B. cereus* can synthesize important molecules for plant growth (Wagi & Ahmed, 2019). When non-rhizobia strains are combined with rhizobia, we predict that these microbes receive a benefit from living inside a nodule because in nature many rhizobia strains will compete for access to nodules where carbon and shelter are provided (Burghardt, 2020). Plants inoculated with both a non-rhizobia and rhizobia strain, however, grew to similar sizes as plants inoculated with just rhizobia suggesting that the non-rhizobia strains do not provide a benefit or cost to plants. We note that these plants were grown in greenhouses under stable conditions, with low nitrogen, and were provided with all other nutrients needed for growth. It is possible that these non-rhizobia strains offer some benefit or cost under field conditions where plants could be experiencing stressful environmental conditions, pathogens, or low nutrients in the soil. For example, some studies have shown that certain non-rhizobia species are effective at mobilizing phosphorus (Arif et al., 2017). Non-rhizobia strains could also be providing a benefit to plants by taking up space in nodules that could otherwise be occupied by pathogens (Boyle et al., 2021; Wang et al., 2022). Overall, under greenhouse conditions, the non-rhizobia strains sourced from tropical and temperate locations were commensal and there was no signal of a benefit to the plant host. Though it is important to note that we only tested a few non-rhizobia strains that can be easily cultured in the lab which could bias results to observing few effects on plant growth.

## Conclusion

Pairing sequence data from natural microbial populations with manipulative experiments is an effective approach to uncovering latitudinal patterns in symbiotic microbe diversity and understanding the impacts of this diversity on host plants. Overall, we found that legumes in the tropics host a higher diversity of non-rhizobia strains in their nodules. It is still unclear what mechanism plants are using to select non-rhizobia strains for entry into nodules since these strains appear to be neutral for host fitness. Given the lack of concordance between soil and nodule microbial communities, plants in the tropics may be less selective in which strains gain entry into nodules.

Our results have important implications for an open question in ecology and evolution: does mutualism drive patterns of diversity or does existing latitudinally structured diversity influence mutualistic interactions? In a legume-rhizobia system where the plant is also interacting with and hosting non-rhizobia strains, host selection appears to play an important role in structuring diversity patterns found in nodule communities. Latitudinal differences in soil nutrients could be an important factor in determining outcomes of host selection in nutrient acquisition mutualisms (Wu et al., 2021). Future work should focus on whether *C. nictitans* plants are locally adapted to their soil environment and microbes or if host selection is plastic.

## Acknowledgements

We would like to thank Stephen Wright, Ben Gilbert, Art Weis, and Shelby Riskin for their feedback on the research. Bill Cole and Tom Gludovacz provided excellent greenhouse support for the experiments. Thank you to Mountain Lake Biological Station, Archbold Biological Station, and the Organization of Tropical Studies for providing lodging and assistance during collection trips. Bonnie Pilkington, Georgia Henry, and Rodolfo Quiros Flores provided invaluable help in the field collecting samples. Oxana Pogouste provided lab support and performed DNA extractions on the soil samples. We also thank the National Commission on Biodiversity Management (Comisión Nacional de Gestión de la Biodiversidad, CONAGEBIO) for granting research permits for collections of molecular samples in Costa Rica (Permit ID: R-051-2017-OT-CONAGEBIO, R-012-2018-OT-CONAGEBIO). Our work is supported by NSERC and Queen Elizabeth Graduate Scholarships (TLH) and NSERC Discovery Grants (JRS, MEF).

## Data Accessibility

All raw sequences will be deposited to Sequence Read Archive database in GenBank at the time of publication. Metadata and code for analyses will be made available on a public repository on GitHub.

## Benefits Generated

Research was completed with the support and assistance of the OTS (Organization of Tropical Studies). Contributions of all individuals including those stationed at OTS research facilities are described in the Acknowledgements. Results of the research have been shared with scientists at the OTS Las Cruces Biological Station through an oral presentation as part of the LSAMP REU (Louis Stokes Alliances for Minority Participation Research Experience for Undergraduates) summer program in 2019. A report of the results has also been submitted to CONAGEBIO (Comisión Nacional para la Gestión de la Biodiversidad) to be shared with the Costa Rican government in accordance with the sampling permit guidelines (permit numbers listed in Acknowledgements). Results of the research have also been shared with the broader scientific community through online presentations at virtual conferences in 2021 (Canadian Society for Ecology and Evolution, Society for the Study of Evolution).

## Author contributions

TLH, JRS, and MEF conceived of the idea and design of the study. TLH collected the samples (with minor assistance from JRS at La Selva); TLH performed the molecular lab work, performed the tropical greenhouse experiment, and analyzed the sequence and experiment data. ZAP performed the temperate greenhouse experiment. TLH, MEF, and JRS interpreted the results and wrote the manuscript.

## Notes

### Competing Interest Statement

The authors have declared no competing interest.

