## Supplemental Materials for "Is there a latitudinal diversity gradient for symbiotic microbes? A case study with sensitive partridge peas"

### Materials and Methods

**Generation of cultured microbe samples.** We used the supernatant created from crushing nodules as starting material for our cultured samples. We pipetted 400 ml of the supernatant into liquid YM media and incubated the samples at 30°C and 200 rpm for 4 days. Cultured samples were stored frozen at -20°C until DNA extraction. We extracted large amounts of DNA (average of 20.8 ng/ul per sample) from the resulting culture according to the instructions in the DNeasy UltraClean Microbial Kit from Qiagen. We obtained 157 cultured samples.

**Analysis for detecting differences in culture and field sample communities.** We analyzed 77 samples in our dataset that had a paired culture and field sample to identify differences in the microbial community caused by our sampling methods. To visualize the differences in samples, we merged ASVs by sample type (field and culture), aggregated ASVs by family, and plotted the relative proportions of the top 10 most common families in the field and culture samples to identify which bacterial families dominate each sample type. We identified the most common genera in field samples and calculated the frequency of these genera observed in the cultured samples and vice versa.

We estimated several beta diversity indices on the field and culture ASV data using the phyloseq package (McMurdie and Holmes 2013). To evaluate similarities in community structure between sample types we estimated Jaccard's distance on the presence/absence ASV data and estimated Bray-Curtis distance on abundance ASV data. We also calculated unweighted unifrac distance to take into consideration phylogenetic differences between ASVs and identify changes in unique variants across sample types. In addition, we estimated weighted unifrac distance to identify differences in common variants between the sample types. We performed principal coordinate analysis (PCoA) on all beta diversity indices and created a scree plot to examine how much variation was summarized on each axis. In all cases, we plotted the first two PCoA axes as the majority of variation (Axis1: 20.6-91.2%, Axis 2: 5.3-15.2%) was along these two axes. To test whether culture and field samples have different microbial communities we calculated the correlation between PCoA axis 1 scores of field and culture samples (for all distance metrics) using Spearman's rho because distributions of axis 1 scores were non-normal.

**Effects of culturing methods on microbial communities.** The field microbial community consisted mostly of bacteria belonging to the Bradyrhizobiaceae family while the most abundant family in the cultured community was the Bacillaceae family (Supp. Fig. 2, Supp Table 1) Most of the ASVs in the Bradyrhizobiaceae family are rhizobia species belonging to the *Bradyrhizobium* genus which contains many species of rhizobia known to form nodules on plant roots. Species belonging to the Bacillaceae family generally lack nod genes in the genome and are unable to form nodules. In general, each field sample contained more ASVs total than the culture samples with field samples containing an average of 16.047 ± 1 s.e. ASVs per sample and culture samples containing an average of 12.259 ASVs per sample. The most common ASV in the field samples belonged to the genus *Bradyrhizobium* and occurred in 49.49% of field samples and in only 2.60% of culture samples. In contrast, the most common ASV in the culture samples was a *Bacillus cereus* strain and occurred in 49.94% of culture samples and was absent from all field samples (Supp. Table 2). When we merged ASVs based on taxonomic identification, the most abundant genus in the culture samples was *Bacillus* (77,184 reads total). The *Bacillus* genus had a total of 28 reads in the field samples. The *Bradyrhizobium* genus was the most abundant genus in field samples (128,282 reads) and fairly abundant in the culture samples (35,475 reads).

The community structure of field and culture samples differed considerably in ASV composition (Supp. Fig. 1.a, 1.b). PCoA plots based on unifrac distance did not show as much separation between the two sample types on axis 1 (25.9% variation explained) indicating that the sample types do not differ as much in their composition of unique variants (Supp. Fig. 1.c, 1.d). The weighted unifrac distance PCoA plot showed clear separation of the field and culture samples on axis 1 (91.2%) indicating that the sample types differ considerably in their composition of common variants (Supp. Fig. 1.d).

We found a non-significant correlation between PCoA axis 1 values of the field and culture samples for Jaccard's and Bray-Curtis distance (Supp. Table 2). Although we found a significant correlation between the sample types for unifrac and weighted unifrac distances indicating that the sample types have phylogenetically similar communities, the Spearman's rho value indicated weak correlation (Supp. Table 2).

Overall, we found that traditional lab culturing methods greatly changed the composition of individual microbial strains in our samples which could bias overall conclusions made about natural microbial communities. Field and culture communities differed significantly in overall community composition and in common variants. The two types of samples were similar in terms of unique variants and overall phylogenetic similarity. Cultured samples were dominated by fast growing non-rhizobia and non-nodulating strains such as *Bacillus* species. In contrast, slow growing rhizobia strains such as *Bradyrhizobium* species were in high abundance in field samples. Likely when a mix of strains are grown in a small amount of liquid culture in the lab, fast growing strains outcompete the slow growing rhizobia strains. We determined that culturing natural microbial communities was not an appropriate method to characterize microbial communities in nodules collected from the field and is likely more of a concern than PCR bias among our strains in MiSeq sequencing. Methods of extracting DNA that do not require any growth phase in the lab show more promise for painting an accurate picture of diversity in natural microbial communities, even though these methods often produce lower yields of DNA. The amplification phase in 16S sequencing methods seems to overcome any problems with low DNA concentration and still produces high quality sequences for analysis.

**Number of nodules.** To explore whether variation in the number of nodules sampled affected diversity metrics, we included nodule number as a covariate in the models and achieved similar results to when the variable was excluded. We also subsampled our dataset for plants that had between 5-10 nodules and again found similar results suggesting that differences in nodule number sampled among plants does not affect bacterial diversity. Therefore, we only report results from the full ASV dataset without including nodule number as a covariate in our models.

**Methods of identifying rhizobia ASVs.** We performed a BLAST search (blastn) on all ASVs that were unidentified at the genus level in our dataset. If one of the top hits in our blastn results belonged to one of the genera in the list of known rhizobia genera, we identified it as a rhizobia ASV.

**PCR identification of strains for co-inoculation experiments.** We isolated pure single microbial strains from nodules from a single temperate plant (latitude 36.5161, longitude -78.7303) and a single tropical plant (latitude 10.3583, longitude -85.3203). We rehydrated and sterilized these field-collected nodules using the same method described in our DNA extraction protocol. We sterilely crushed each nodule in a small amount of water in a 1.5 ml Eppendorf tube. We plated the liquid onto YM agar plates with sterile plastic inoculation loops and incubated the plates at 30°C for up to 10 days or until sufficient colony growth was observed. We chose one colony on the plate to re-streak on a fresh plate. We repeated the re-streaking process three times to isolate a single pure strain. We then inoculated 7 ml of liquid YM media with the pure colony and incubated the sample in a shaker at 30°C and 200 rpm for four days. We froze 1 ml of the culture mixed with 50% glycerol solution in liquid nitrogen for storage in a -80°C freezer. We performed DNA extractions on 2 ml of culture using the DNeasy UltraClean Microbial Kit from Qiagen. We performed polymerase chain reactions (PCR) on the DNA samples to amplify the 16S rDNA locus using 16S_27F (AGAGTTTGATCMTGGCTCAG) and 16S_1492R primers (GGTTACCTTGTTACGACTT) and PCR conditions outlined in (Weisburg et al. 1991). We submitted the 16S amplicons for Sanger sequencing at CAGEF at the University of Toronto and analyzed the sequences in Geneious (version 9.1.8). We performed a BLAST search (blastn with default parameters) on the consensus sequences against Genbank to identify the species of microbes. We identified 3 unique species associated with the temperate plant: one rhizobia species *Bradyrhizobium elkanii* and two non-rhizobia species *Pseudomonas korensis* and *Variovorax paradoxus.* We identified one rhizobia species associated with the tropical plant *Bradyrhizobium yuanmingense* and one non-rhizobia species *Bacillus cereus*.

**Growth chamber and greenhouse conditions.** Growth spaces were sterilized before running experiments by heating the spaces at 35°C for two days and sanitizing surfaces with a spray consisting of peracetic acid and hydrogen peroxide. For both experiments, we prepared seeds by scarifying them with sandpaper and then sterilizing them in 95% ethanol for 20 seconds and then in bleach for 20 seconds. We rinsed seeds with sterile water and stratified seeds on moist filter paper in a sealed petri dish at 30°C overnight. Germinated seeds were planted in sterile sand in Magenta Boxes (tropical experiment) or in autoclavable Cone-tainers (temperate experiment). We prepared Magenta Boxes and Cone-tainers ahead of time by packing with sand, inserting a polypropylene wick, and autoclaving twice on a one hour 121°C cycle. The Cone-tainers were plugged with cotton balls to hold sand in place. Sterile 15 ml Falcon tubes were used to hold water under the Cone-tainers and connect to the wick. The bottom compartments of the Magenta Boxes and the Falcon tubes were filled with an autoclaved low-nitrogen Fahräeus fertilizer (Zhang et al. 2020) to supply plants with enough nutrients for growth. Every other week the fertilizer was replenished and in the alternate weeks the bottoms were filled with autoclaved distilled water. Growth chamber conditions were set at a day temperature of 26-27 °C, a night temperature of 18-19 °C, and a light period of 15 hours. Microbial cultures were created by thawing the frozen microbial sample and plating the solution on a sterile YMA plate. After re-streaking the plate three times we inoculated liquid YM media with the cultures and grew the solution to an OD600 reading of either 0.2 (for the tropical experiment) or 0.4 (temperate experiment) at 30°C and 200 rpm. We inoculated each plant with 2 ml of culture once in the temperate experiment and twice in the tropical experiment (once a week after planting and again at three weeks after planting).

### Figures

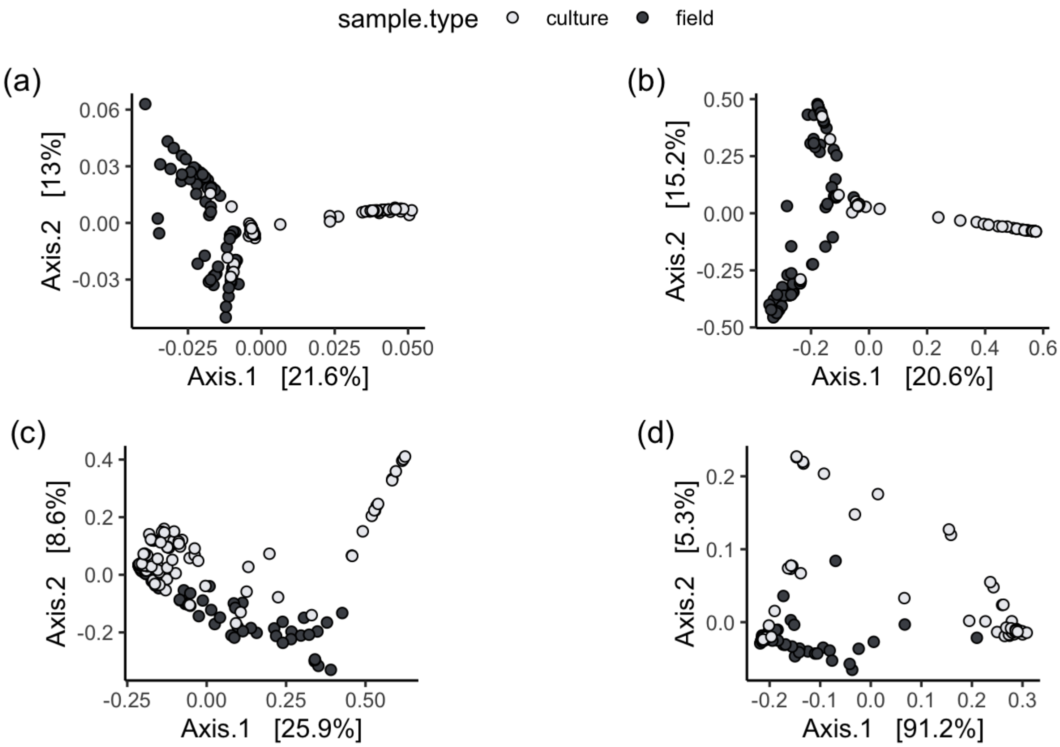


**Supplemental Figure 1. PCoA plots of cultured (light grey) and field (dark grey) sampled nodule microbial communities.** Different plots show results from the different distance metrics performed on the data (a) Jaccard's distance, (b) Bray-Curtis, (c) unifrac distance, (d) weighted unifrac distance.

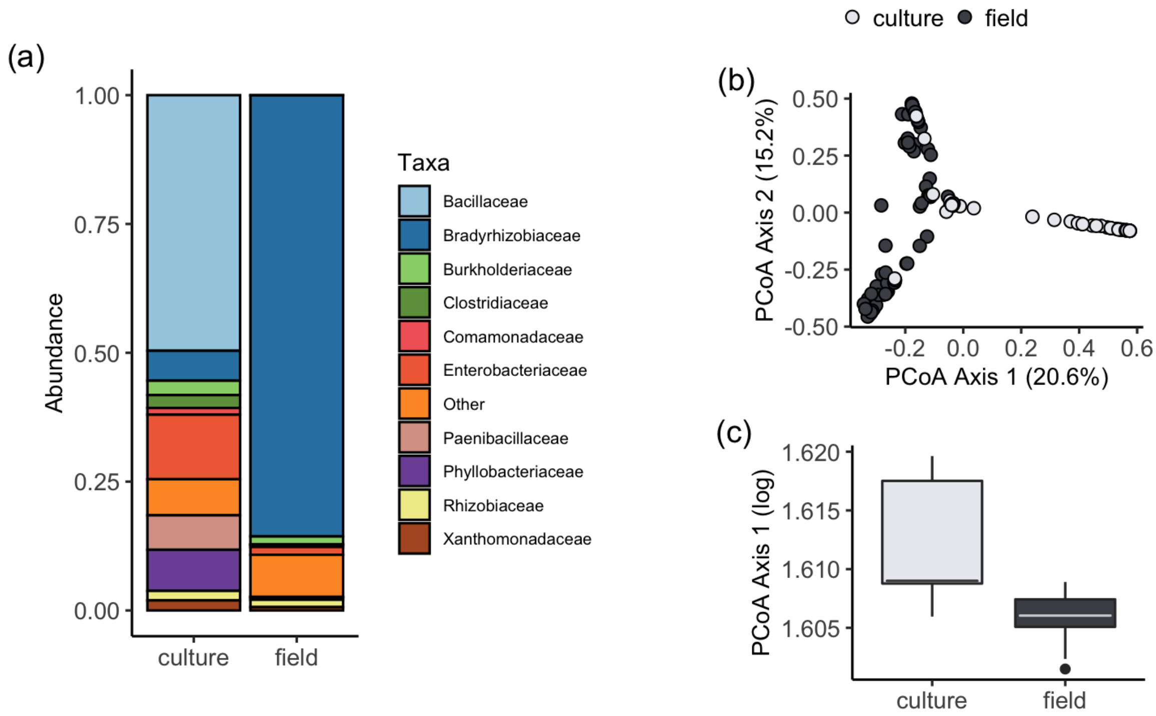


**Supplemental Figure 2. Breakdown of the top 10 most common taxonomic families of microbes found in culture samples and field samples.** Families were identified on ASVs from 16S sequence data using the Greengenes database. Culture samples were grown to high OD600 readings in the lab before sequencing. Field samples were sequenced directly from the supernatant of crushed nodules.

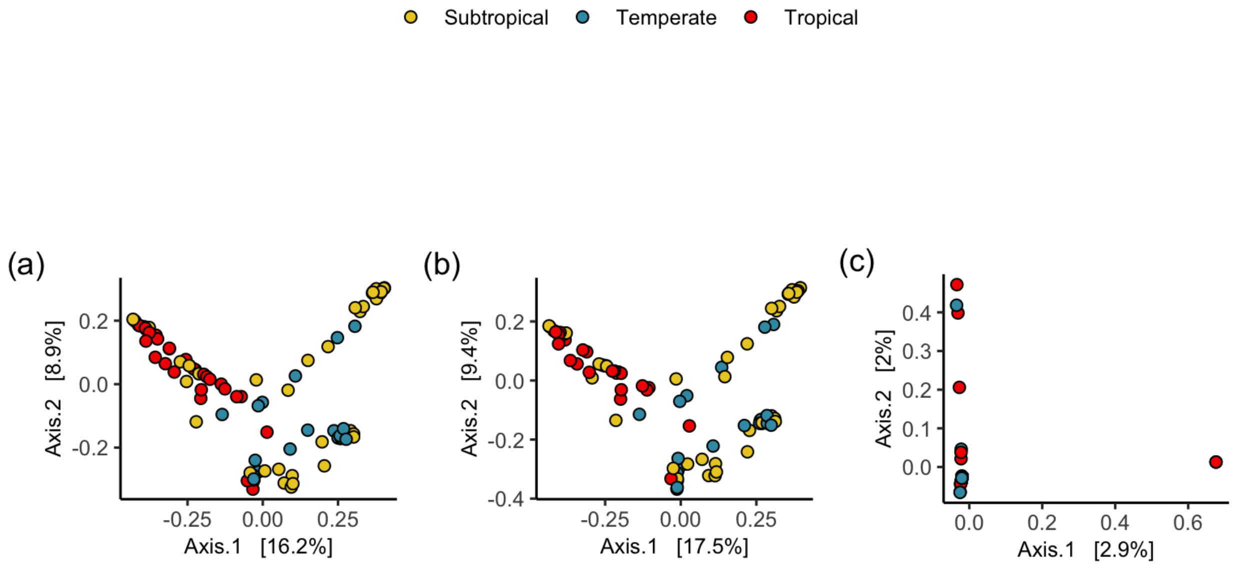


**Supplemental Figure 3. Principal Coordinate Analysis (PCoA) of community structure based on Jaccard's dissimilarity in nodule communities from *Chamaecrista nictitans* nodules across latitude.** Each dot represents a single plant where nodules were pooled and the microbe community was extracted and sequenced. Panel (a) represents PCoA performed on the full ASV community. The middle panel (b) represents rhizobia only ASVs and the last column (c) are non-rhizobia ASVs.

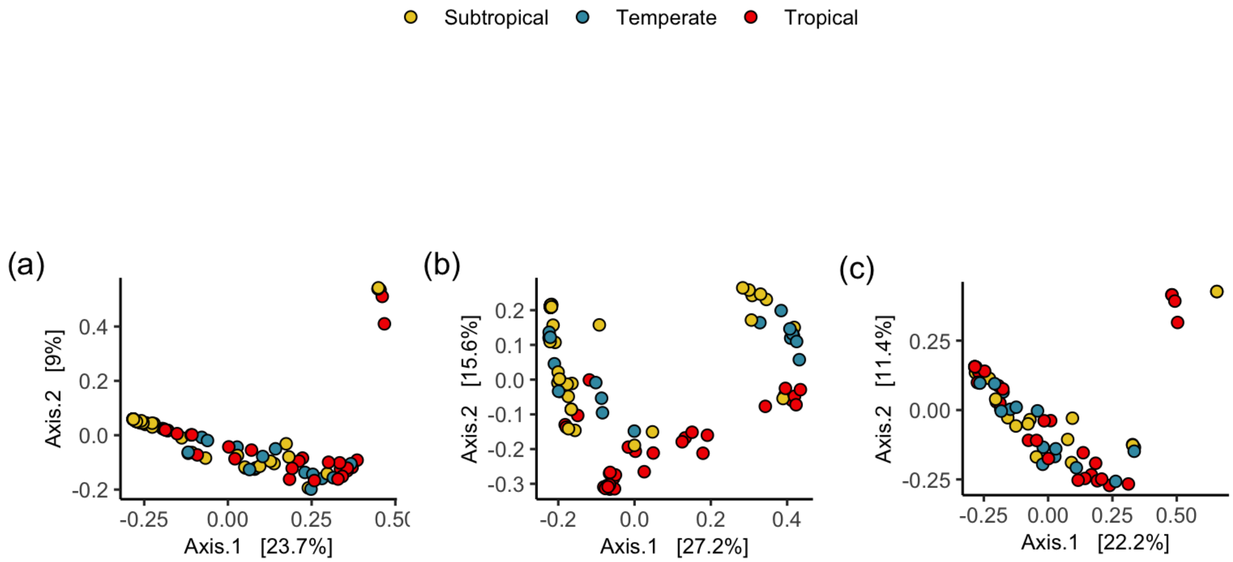


**Supplemental Figure 4. Principal Coordinate Analysis (PCoA) of community structure based on Unifrac dissimilarity in nodule communities from *Chamaecrista nictitans* nodules across latitude.** Each dot represents a single plant where nodules were pooled and the microbe community was extracted and sequenced. Panel (a) represents PCoA performed on the full ASV community. The middle panels (b) represent rhizobia only ASVs and the last column (c) are non-rhizobia ASVs.

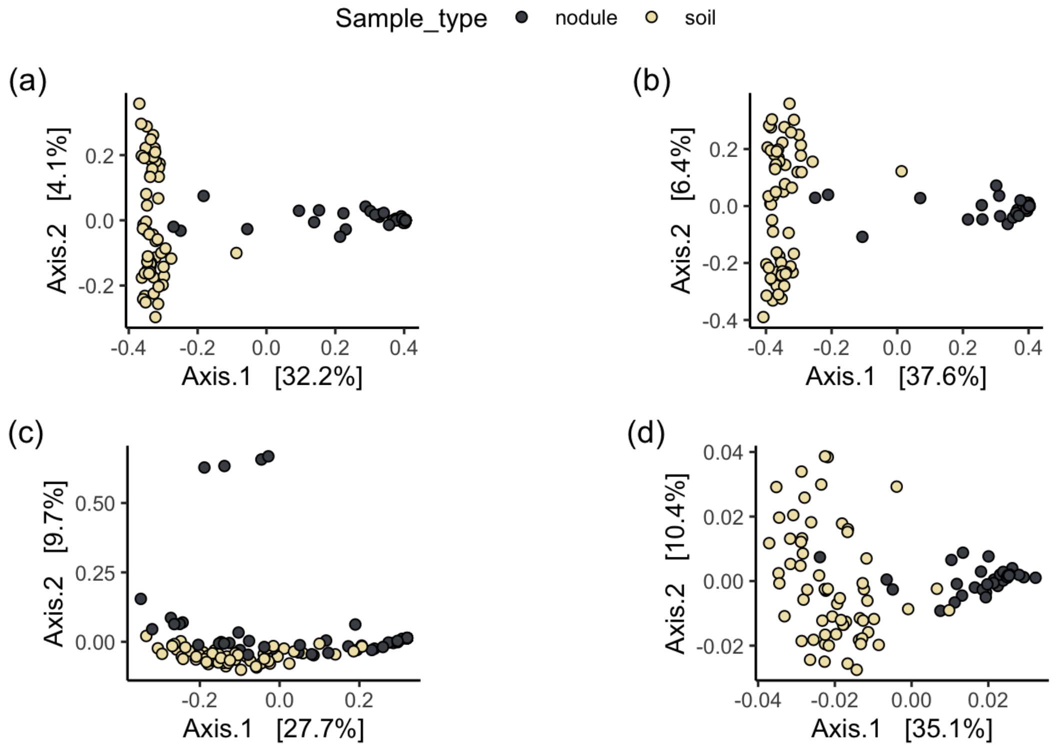


**Supplemental Figure 5. PCoA plots of nodule isolated microbes (dark grey) and soil community microbes (light yellow).** Soil microbes represent ASVs merged to 97% similarity and also found in nodules. Different plots show results from the different distance metrics performed on the data (a) Jaccard's distance, (b) Bray-Curtis, (c) unifrac distance, (d) weighted unifrac distance.

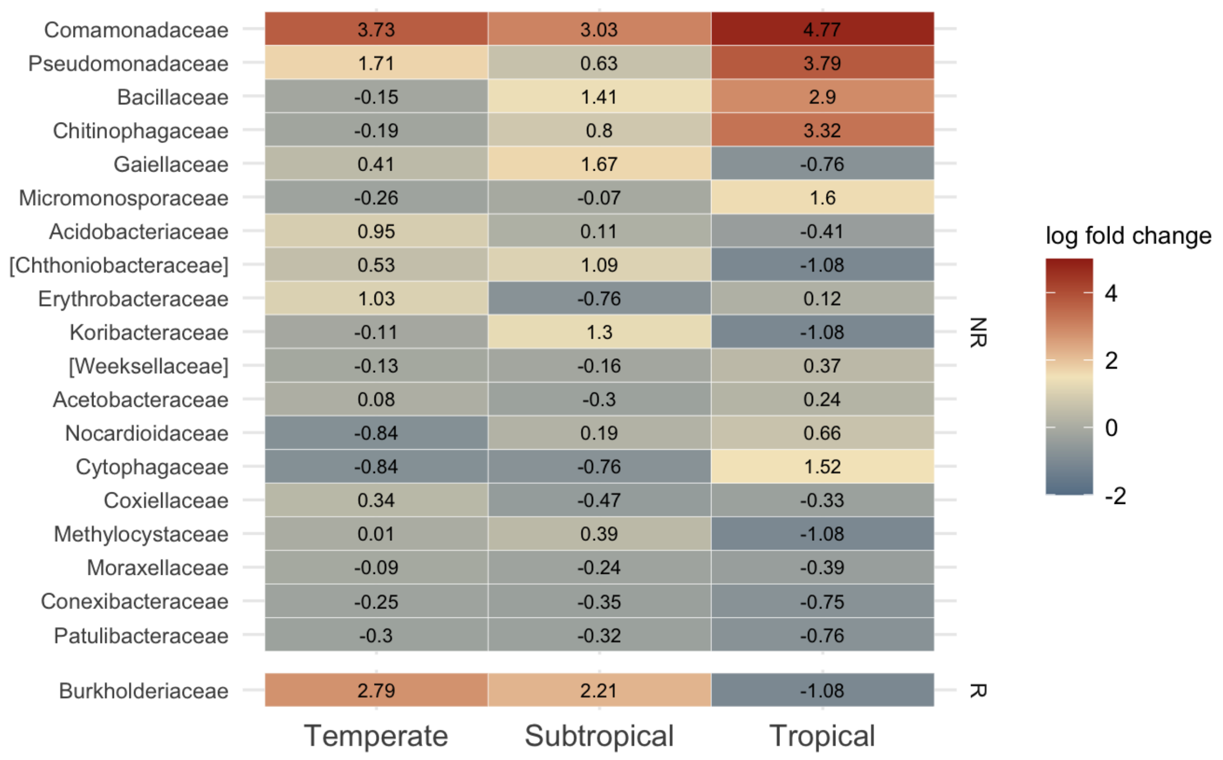


**Supplemental Figure 6. Heat map representing the log fold change in abundance of different microbial genera in the temperate, subtropical, and tropical soils**. Differential abundance analysis with ANCOMBC was performed on non-rarefied raw OTU data where ASVs were merged by 97% similarity. Soil OTUs were subset to OTUs that could occupy nodules prior to analysis. R represents rhizobia families and NR represents non-rhizobia families. Red represents genera with high abundance in the samples and blue represents genera with low abundance. All families plotted showed significance in the differential abundance analysis.

### Tables

**Supplemental Table 1. Most common ASVs in field and culture samples.** Prevalence reported here is the proportion of plant samples that ASV is found in. Each sample has a culture community and field community. Total of 77 samples. ASVs reported here are the rarefied ASVs calculated from unique exact sequences.

| ASV code | taxonomic id | field prevalence | culture prevalence |
|---|---|---|---|
| *top culture ASVs* | | | |
| ASV1 | *Bacillus cereus* | 0 | 0.494 |
| ASV2 | *Bacillus cereus* | 0 | 0.494 |
| ASV3 | *Bacillus cereus* | 0 | 0.481 |
| ASV4 | *Bacillus cereus* | 0 | 0.468 |
| ASV5 | *Bacillus cereus* | 0 | 0.468 |
| *top field ASVs* | | | |
| ASV6 | *Bradyrhizobium* | 0.494 | 0.026 |
| ASV7 | *Bradyrhizobium* | 0.481 | 0.026 |
| ASV8 | *Bradyrhizobium* | 0.455 | 0.026 |
| ASV9 | *Bradyrhizobium* | 0.429 | 0.013 |
| ASV10 | *Bradyrhizobiaceae* | 0.429 | 0.052 |

**Supplemental Table 2. Results of culture PCoA axis 1 scores regressed on field PCoA axis 1 scores for different beta diversity metrics in linear models.** ASVs used in the analysis are exact sequences. Wald $\chi^2$ values and p values are the results of type III Anova tests. Population was included as a random effect in the models. When necessary, we squared or calculated the inverse of the response variable to fit model assumptions.

| **field axis1** | **estimate** | **s.d.** | **Wald $\chi^2$** | **d.f.** | **p value** |
|---|---|---|---|---|---|
| *ASV, n = 76* | | | | | |
| Jaccard | -0.0157 | 0.0135 | 1.3584 | 1 | 0.2438 |
| Bray Curtis | 0.052 | 0.2203 | 0.0553 | 1 | 0.8141 |
| Unifrac | -20.635 | 16.316 | 1.5996 | 1 | 0.2060 |
| Weighted unifrac | 0.0120 | 0.0351 | 0.1168 | 1 | 0.7325 |

**Supplemental Table 3. Differences in the most common ASVs found in tropical, subtropical, and temperate zones.** Values represent the proportion of samples within each region that particular ASV is found in. ASVs are exact unique sequences of the 16S locus.

| ASV code | taxonomic id | tropical | subtropical | temperate |
|---|---|---|---|---|
| *top tropical ASVs* | | | | |
| ASV1 | *Bradyrhizobium* | 0.844 | 0.250 | 0 |
| ASV2 | *Bradyrhizobium* | 0.844 | 0.295 | 0.059 |
| ASV3 | *Bradyrhizobium* | 0.844 | 0.273 | 0.059 |
| ASV4 | *Bradyrhizobium* | 0.781 | 0.227 | 0.059 |
| ASV5 | *Bradyrhizobium* | 0.688 | 0.159 | 0.059 |
| *top subtropical ASVs* | | | | |
| ASV6 | *Bradyrhizobium* | 0.031 | 0. 773 | 0.765 |
| ASV7 | *Bradyrhizobium* | 0.031 | 0. 773 | 0.706 |
| ASV8 | *Bradyrhizobium* | 0.031 | 0. 737 | 0.647 |
| ASV9 | *Bradyrhizobium* | 0 | 0. 591 | 0.529 |
| ASV10 | *Bradyrhizobium* | 0.031 | 0. 591 | 0.588 |
| *top temperate ASVs* | | | | |
| ASV6 | *Bradyrhizobium* | 0.031 | 0.773 | 0.765 |
| ASV7 | *Bradyrhizobium* | 0.031 | 0.773 | 0.706 |
| ASV8 | *Bradyrhizobium* | 0.031 | 0.737 | 0.647 |
| ASV10 | *Bradyrhizobium* | 0.031 | 0.591 | 0.588 |
| ASV9 | *Bradyrhizobium* | 0 | 0.591 | 0.529 |

**Supplemental Table 4. Results of nodule PCoA axis 1 scores regressed on soil PCoA axis 1 scores for different beta diversity metrics in linear models.** Soil OTUs used in the analysis are filtered to OTUs shared between nodules and soil samples. Wald $\chi^2$ values and p values are the results of type III Anova tests. Population was included as a random effect in the models.

| **soil axis1** | **estimate** | **s.d.** | **Wald $\chi^2$** | **d.f.** | **p value** |
|---|---|---|---|---|---|
| *OTUs found in nodules, n = 61* | | | | | |
| Jaccard | -0.3019 | 0.5024 | 0.3612 | 1 | 0.5479 |
| Bray Curtis | -0.3198 | 0.2847 | 1.2617 | 1 | 0.2613 |
| Unifrac | 0.0994 | 0.2151 | 0.1971 | 1 | 0.6571 |
| Weighted unifrac | -0.0987 | 0.1121 | 0.7748 | 1 | 0.3787 |

**Supplemental Table 5. Results of linear models testing the impact of latitude of sample on the community difference between nodule and soil microbes.** Soil OTUs used in the analysis are filtered to OTUs shared between nodules and soil samples. Bolded factors represent significance at p<0.05.

| | estimate | s.d. | F | d.f. | p value |
|---|---|---|---|---|---|
| *OTUs found in nodules, n = 61* | | | | | |
| Jaccard | -0.0002 | 0.0004 | 0.1647 | 1 | 0.6864 |
| Bray Curtis | -0.0004 | 0.0013 | 0.0007 | 1 | 0.7519 |
| **Unifrac** | **-0.0060** | **0.0012** | **25.5070** | **1** | **<0.0001** |
| Weighted unifrac | -0.0002 | 0.0002 | 1.6025 | 1 | 0.2105 |
